## Supplementary Figures for "OpenSpliceAI: An efficient, modular implementation of SpliceAI enabling easy retraining on non-human species"

[†]These authors contributed equally

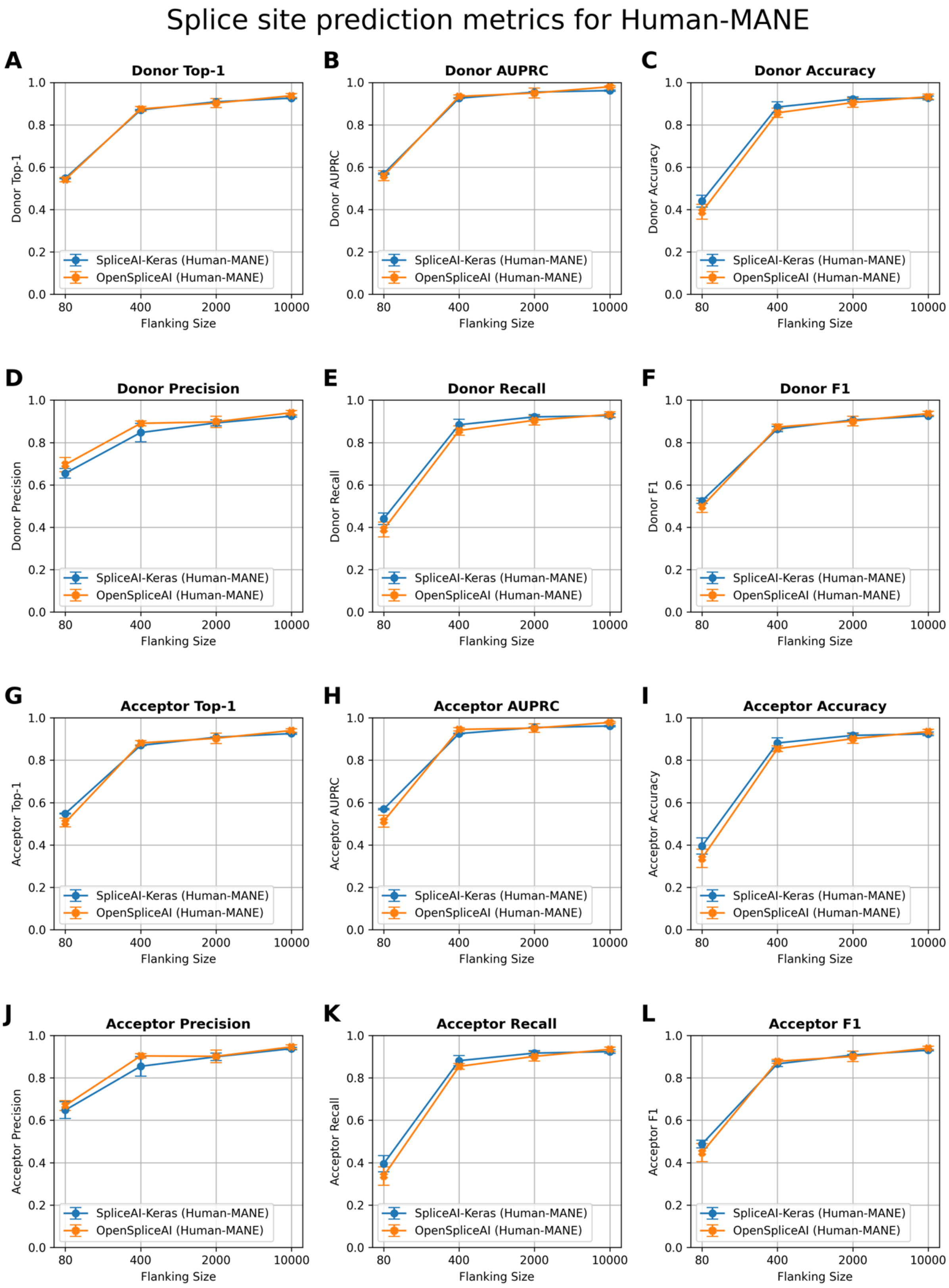


**Figure S1.** Comparison of splice site prediction performance between SpliceAI-Keras (blue) and OSAI$_{MANE}$ (orange) across human (*Homo sapiens*) datasets with varying flanking sequence lengths. The plots display donor (5′) and acceptor (3′) splice site prediction metrics using 80, 400, 2000, and 10,000 nt of flanking context. $OSAI_{MANE}$ was trained using the RefSeq MANE v1.3 database(https://ftp.ncbi.nlm.nih.gov/refseq/MANE/MANE_human/release_1.3/MANE.GRCh38.v1.3.refseq_genomic.gff.gz) and GRCh38.p14 genome. **(A)** Donor Top-1: measures the percentage of times the model's most confident prediction exactly matches the true donor site label. **(B)** Donor AUPRC: Area Under the Precision–Recall Curve for donor site predictions. **(C)** Donor Accuracy: proportion of correct donor site calls among all predictions. **(D)** Donor Precision: the fraction of predicted donor sites that are correct. **(E)** Donor Recall: the fraction of true donor sites that are correctly predicted. **(F)** Donor F1: the harmonic mean of precision and recall for donor sites. **(G)** Acceptor Top-1: measures the percentage of times the model's most confident prediction exactly matches the true acceptor site label. **(H)** Acceptor AUPRC: Area Under the Precision–Recall Curve for acceptor site predictions. **(I)** Acceptor Accuracy: proportion of correct acceptor site calls among all predictions. **(J)** Acceptor Precision: the fraction of predicted acceptor sites that are correct. **(K)** Acceptor Recall: the fraction of true acceptor sites that are correctly predicted. **(L)** Acceptor F1: the harmonic mean of precision and recall for acceptor sites. Each panel illustrates that increasing the flanking sequence length generally enhances model performance, with both SpliceAI-Keras and OpenSpliceAI achieving high accuracy and F1 scores at 10,000 nt. Each data point represents the mean across five independently trained models, with error bars indicating the standard deviation.

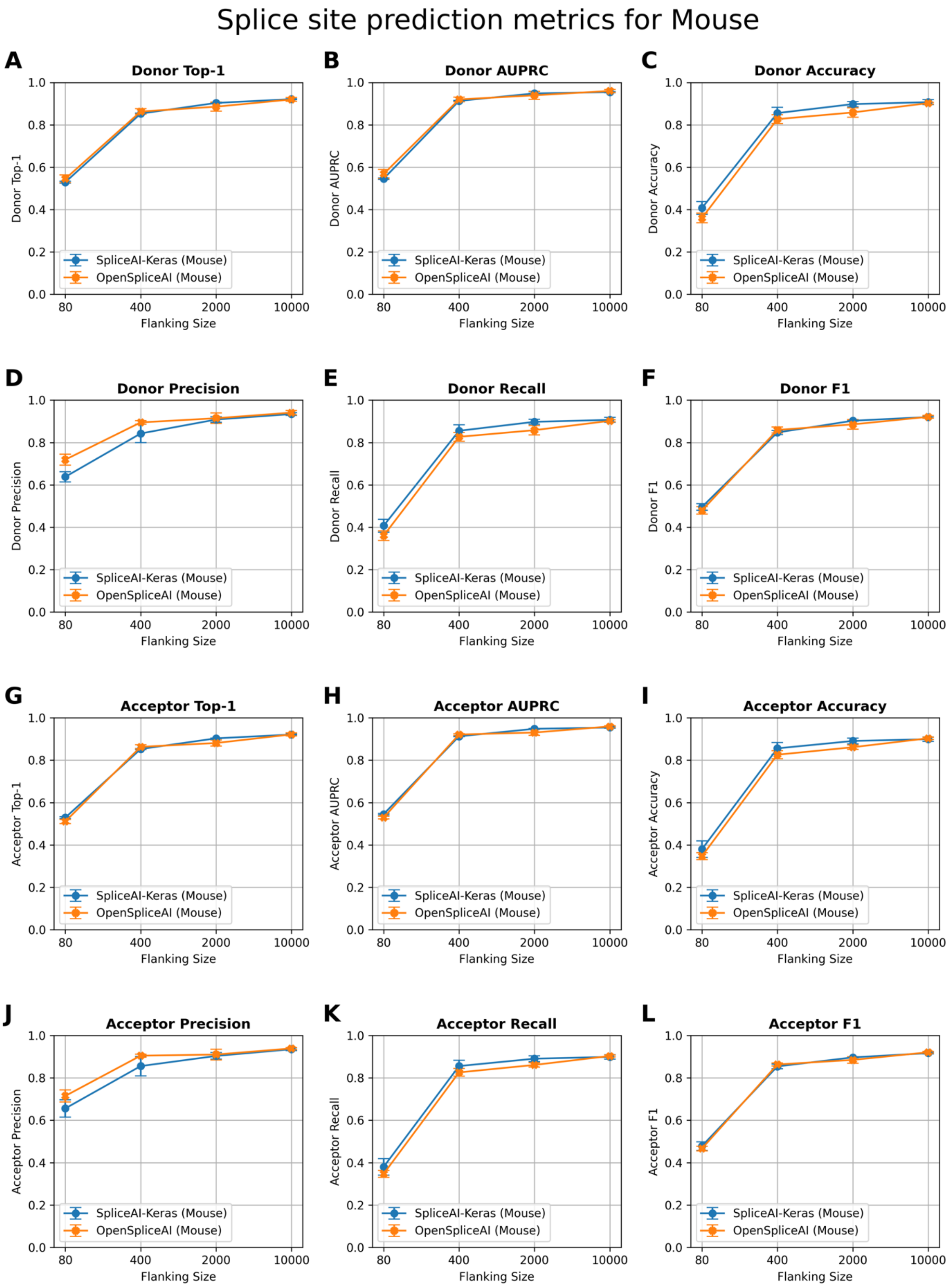


**Figure S2.** Comparison of splice site prediction performance between SpliceAI-Keras (blue) and $OSAI_{MANE}$ (orange) across house mouse (*Mus musculus*) datasets with varying flanking sequence lengths. The plots display donor (5′) and acceptor (3′) splice site prediction metrics using 80, 400, 2000, and 10,000 nt of flanking context. $OSAI_{MANE}$ was trained using the RefSeq MANE v1.3 database(https://ftp.ncbi.nlm.nih.gov/refseq/MANE/MANE_human/release_1.3/MANE.GRCh38.v1.3.refseq_genomic.gff.gz) and GRCh38.p14 genome. The house mouse datasets are curated from RefSeq GRCm39 annotation (https://ftp.ncbi.nlm.nih.gov/genomes/all/GCF/000/001/635/GCF_000001635.27_GRCm39/GCF_000001635.27_GRCm39_genomic.gff.gz) and GRCm39 genome. **(A)** Donor Top-1: measures the percentage of times the model's most confident prediction exactly matches the true donor site label. **(B)** Donor AUPRC: Area Under the Precision–Recall Curve for donor site predictions. **(C)** Donor Accuracy: proportion of correct donor site calls among all predictions. **(D)** Donor Precision: the fraction of predicted donor sites that are correct. **(E)** Donor Recall: the fraction of true donor sites that are correctly predicted. **(F)** Donor F1: the harmonic mean of precision and recall for donor sites. **(G)** Acceptor Top-1: measures the percentage of times the model's most confident prediction exactly matches the true acceptor site label. **(H)** Acceptor AUPRC: Area Under the Precision–Recall Curve for acceptor site predictions. **(I)** Acceptor Accuracy: proportion of correct acceptor site calls among all predictions. **(J)** Acceptor Precision: the fraction of predicted acceptor sites that are correct. **(K)** Acceptor Recall: the fraction of true acceptor sites that are correctly predicted. **(L)** Acceptor F1: the harmonic mean of precision and recall for acceptor sites. Each panel illustrates that increasing the flanking sequence length generally enhances model performance, with both SpliceAI-Keras and OpenSpliceAI achieving high accuracy and F1 scores at 10,000 nt. Each data point represents the mean across five independently trained models, with error bars indicating the standard deviation.

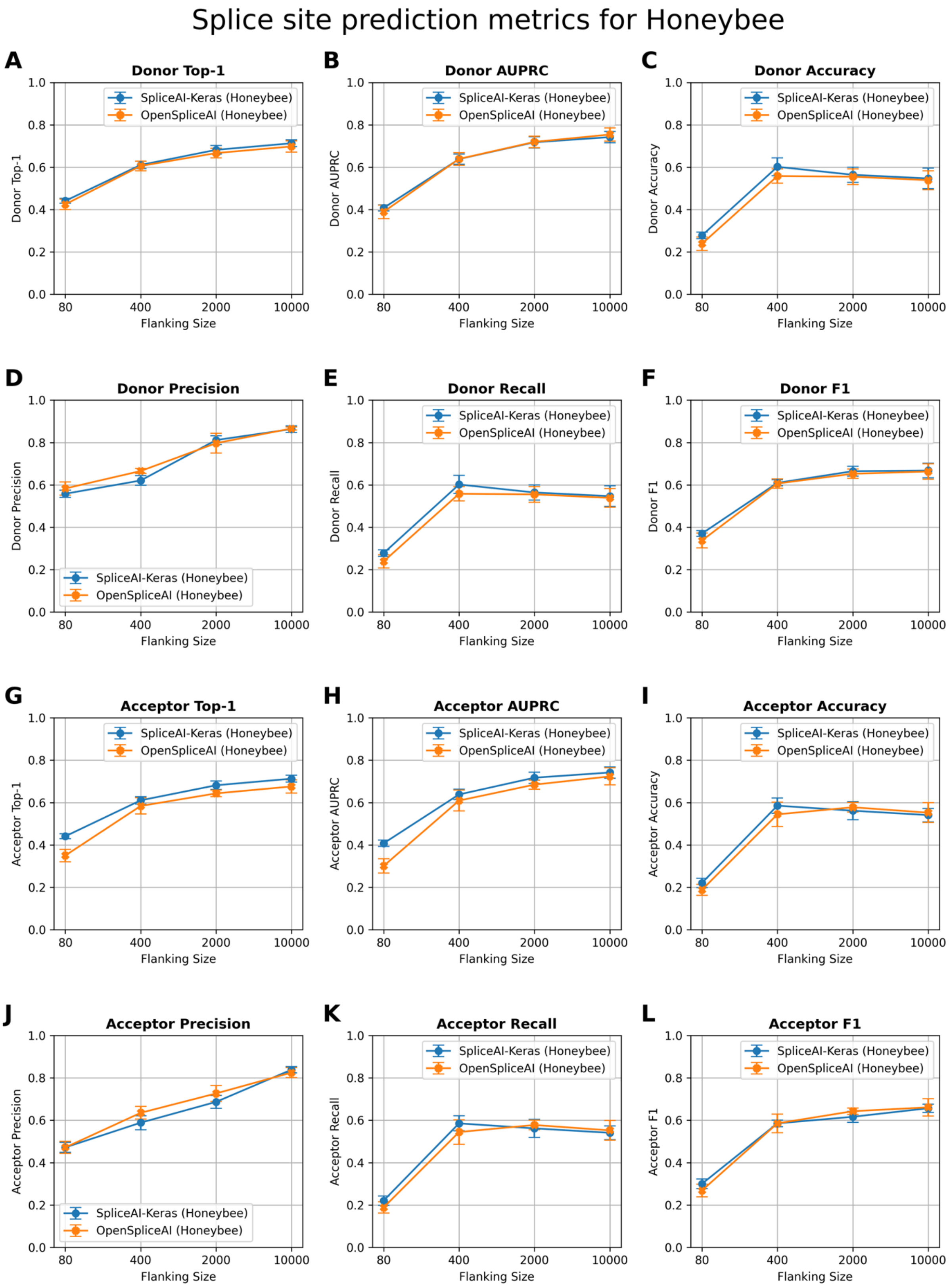


**Figure S3.** Comparison of splice site prediction performance between SpliceAI-Keras (blue) and $OSAI_{MANE}$ (orange) across honeybee (*Apis mellifera*) datasets with varying flanking sequence lengths. The plots display donor (5′) and acceptor (3′) splice site prediction metrics using 80, 400, 2000, and 10,000 nt of flanking context. $OSAI_{MANE}$ was trained using the RefSeq MANE v1.3 database(https://ftp.ncbi.nlm.nih.gov/refseq/MANE/MANE_human/release_1.3/MANE.GRCh38.v1.3.refseq_genomic.gff.gz) and GRCh38.p14 genome. The honeybee datasets are curated from RefSeq Amel_HAv3.1 annotation (https://ftp.ncbi.nlm.nih.gov/genomes/all/GCF/003/254/395/GCF_003254395.2_Amel_HAv3.1/GCF_003254395.2_Amel_HAv3.1_genomic.gff.gz) and Amel_HAv3.1 genome. **(A)** Donor Top-1: measures the percentage of times the model's most confident prediction exactly matches the true donor site label. **(B)** Donor AUPRC: Area Under the Precision–Recall Curve for donor site predictions. **(C)** Donor Accuracy: proportion of correct donor site calls among all predictions. **(D)** Donor Precision: the fraction of predicted donor sites that are correct. **(E)** Donor Recall: the fraction of true donor sites that are correctly predicted. **(F)** Donor F1: the harmonic mean of precision and recall for donor sites. **(G)** Acceptor Top-1: measures the percentage of times the model's most confident prediction exactly matches the true acceptor site label. **(H)** Acceptor AUPRC: Area Under the Precision–Recall Curve for acceptor site predictions. **(I)** Acceptor Accuracy: proportion of correct acceptor site calls among all predictions. **(J)** Acceptor Precision: the fraction of predicted acceptor sites that are correct. **(K)** Acceptor Recall: the fraction of true acceptor sites that are correctly predicted. **(L)** Acceptor F1: the harmonic mean of precision and recall for acceptor sites. Each panel illustrates that increasing the flanking sequence length generally enhances model performance, with both SpliceAI-Keras and OpenSpliceAI achieving high accuracy and F1 scores at 10,000 nt. Each data point represents the mean across five independently trained models, with error bars indicating the standard deviation.

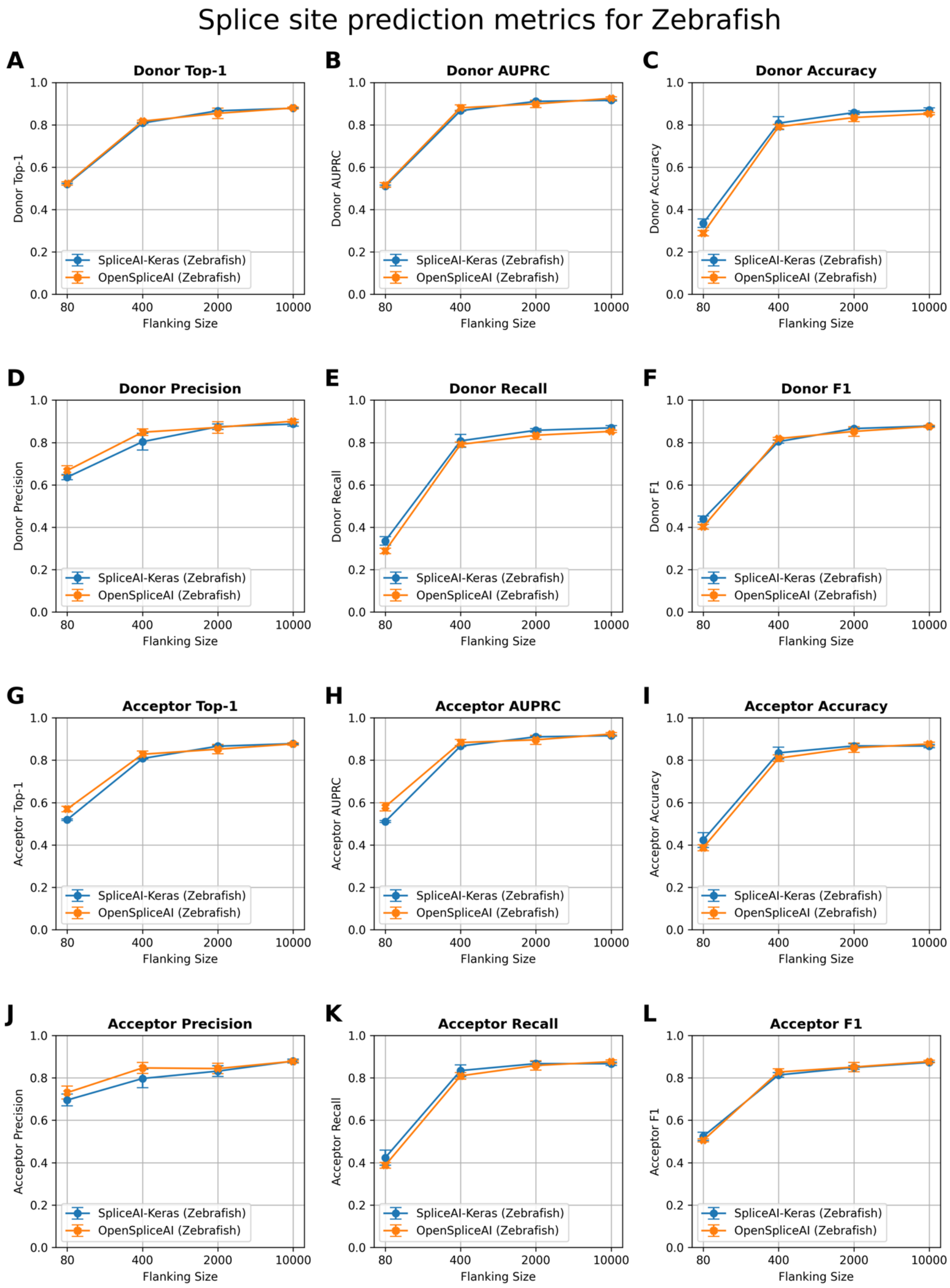


**Figure S4.** Comparison of splice site prediction performance between SpliceAI-Keras (blue) and $OSAI_{MANE}$ (orange) across zebrafish (*Danio rerio*) datasets with varying flanking sequence lengths. The plots display donor (5′) and acceptor (3′) splice site prediction metrics using 80, 400, 2000, and 10,000 nt of flanking context. OSAI$_{MANE}$ was trained using the RefSeq MANE v1.3 database(https://ftp.ncbi.nlm.nih.gov/refseq/MANE/MANE_human/release_1.3/MANE.GRCh38.v1.3.refseq_genomic.gff.gz) and GRCh38.p14 genome. The zebrafish datasets are curated from RefSeq GRCz11 annotation (https://ftp.ncbi.nlm.nih.gov/genomes/all/GCF/000/002/035/GCF_000002035.6_GRCz11/GCF_000002035.6_GRCz11_genomic.gff.gz) and GRCz11 genome. **(A)** Donor Top-1: measures the percentage of times the model's most confident prediction exactly matches the true donor site label. **(B)** Donor AUPRC: Area Under the Precision–Recall Curve for donor site predictions. **(C)** Donor Accuracy: proportion of correct donor site calls among all predictions. **(D)** Donor Precision: the fraction of predicted donor sites that are correct. **(E)** Donor Recall: the fraction of true donor sites that are correctly predicted. **(F)** Donor F1: the harmonic mean of precision and recall for donor sites. **(G)** Acceptor Top-1: measures the percentage of times the model's most confident prediction exactly matches the true acceptor site label. **(H)** Acceptor AUPRC: Area Under the Precision–Recall Curve for acceptor site predictions. **(I)** Acceptor Accuracy: proportion of correct acceptor site calls among all predictions. **(J)** Acceptor Precision: the fraction of predicted acceptor sites that are correct. **(K)** Acceptor Recall: the fraction of true acceptor sites that are correctly predicted. **(L)** Acceptor F1: the harmonic mean of precision and recall for acceptor sites. Each panel illustrates that increasing the flanking sequence length generally enhances model performance, with both SpliceAI-Keras and OpenSpliceAI achieving high accuracy and F1 scores at 10,000 nt. Each data point represents the mean across five independently trained models, with error bars indicating the standard deviation.

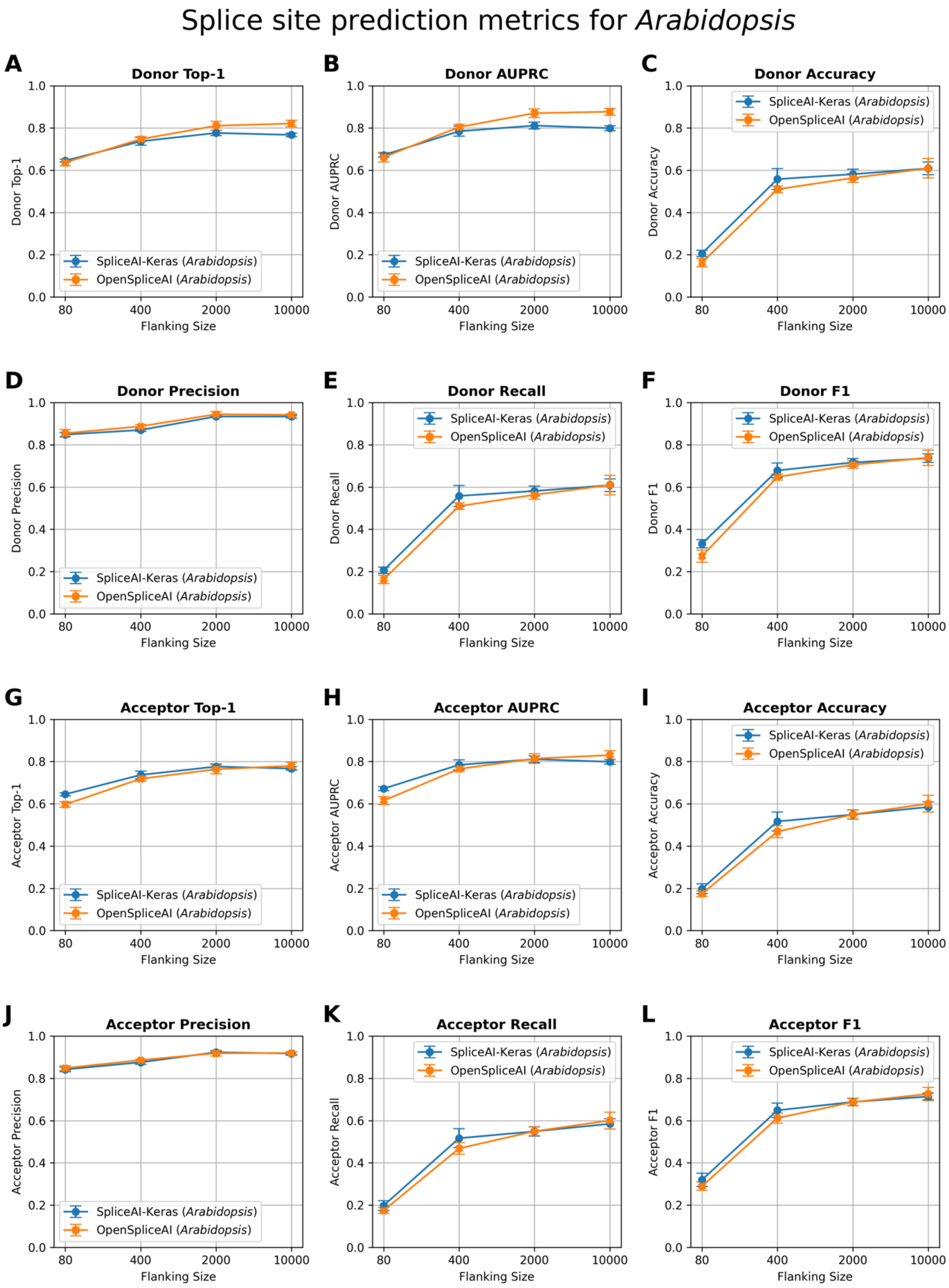


**Figure S5.** Comparison of splice site prediction performance between SpliceAI-Keras (blue) and OSAI$_{MANE}$ (orange) across *Arabidopsis thaliana* datasets with varying flanking sequence lengths. The plots display donor (5′) and acceptor (3′) splice site prediction metrics using 80, 400, 2000, and 10,000 nt of flanking context. $OSAI_{MANE}$ was trained using the RefSeq MANE v1.3 database(https://ftp.ncbi.nlm.nih.gov/refseq/MANE/MANE_human/release_1.3/MANE.GRCh38.v1.3.refseq_genomic.gff.gz) and GRCh38.p14 genome. The *Arabidopsis thaliana* datasets are curated from RefSeq TAIR10.1 annotation (https://ftp.ncbi.nlm.nih.gov/genomes/all/GCF/000/001/735/GCF_000001735.4_TAIR10.1/GCF_000001735.4_TAIR10.1_genomic.gff.gz) and TAIR10.1 genome. **(A)** Donor Top-1: measures the percentage of times the model's most confident prediction exactly matches the true donor site label. **(B)** Donor AUPRC: Area Under the Precision–Recall Curve for donor site predictions. **(C)** Donor Accuracy: proportion of correct donor site calls among all predictions. **(D)** Donor Precision: the fraction of predicted donor sites that are correct. **(E)** Donor Recall: the fraction of true donor sites that are correctly predicted. **(F)** Donor F1: the harmonic mean of precision and recall for donor sites. **(G)** Acceptor Top-1: measures the percentage of times the model's most confident prediction exactly matches the true acceptor site label. **(H)** Acceptor AUPRC: Area Under the Precision–Recall Curve for acceptor site predictions. **(I)** Acceptor Accuracy: proportion of correct acceptor site calls among all predictions. **(J)** Acceptor Precision: the fraction of predicted acceptor sites that are correct. **(K)** Acceptor Recall: the fraction of true acceptor sites that are correctly predicted. **(L)** Acceptor F1: the harmonic mean of precision and recall for acceptor sites. Each panel illustrates that increasing the flanking sequence length generally enhances model performance, with both SpliceAI-Keras and OpenSpliceAI achieving high accuracy and F1 scores at 10,000 nt. Each data point represents the mean across five independently trained models, with error bars indicating the standard deviation.

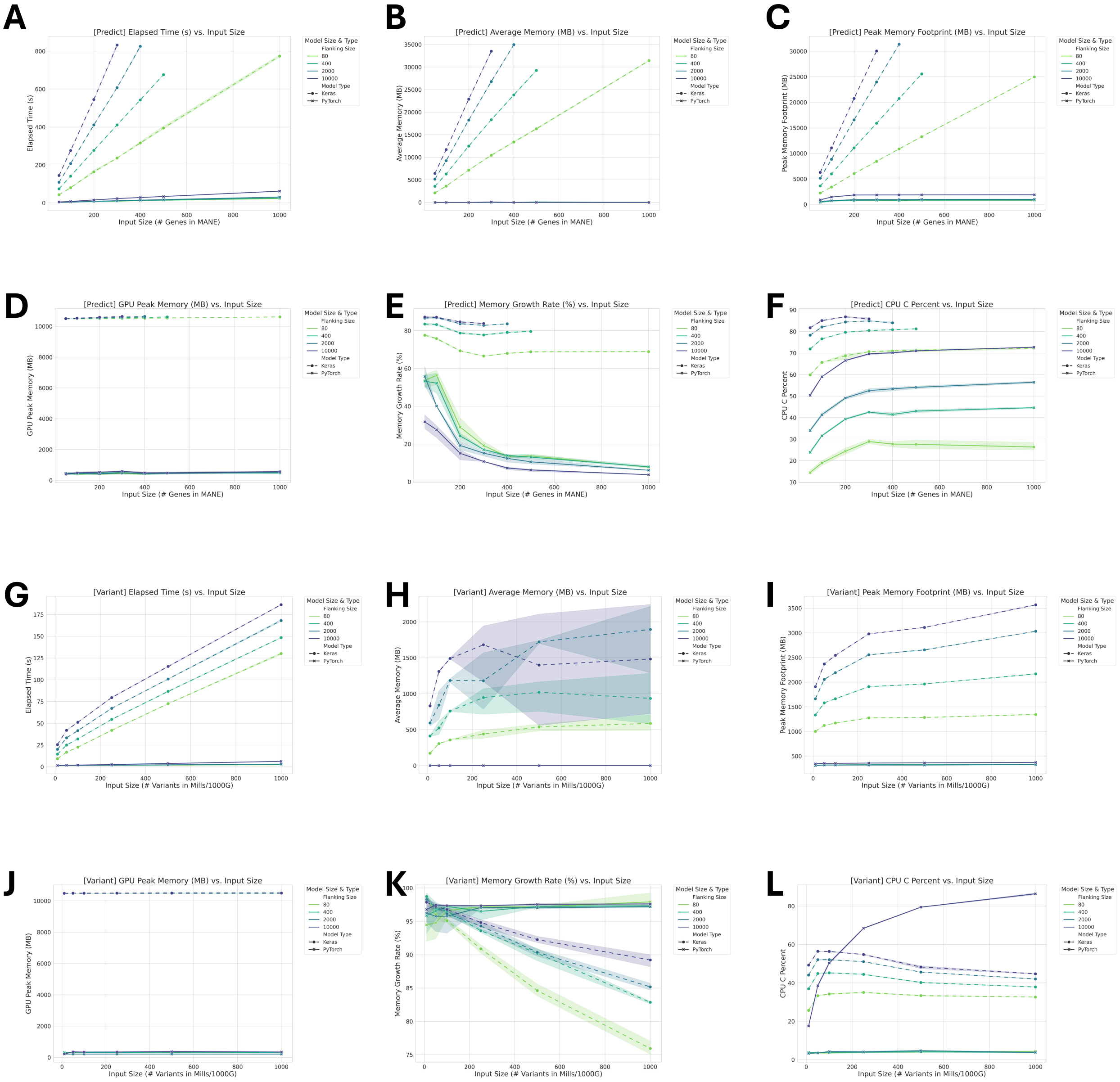


**Figure S6.** Comparison of runtime and memory metrics for `predict` (panels A–F) and `variant` (panels G–L) in OSAI$_{MANE}$ models with different flanking sequences. Each corresponding pair of panels displays the same metric for the two methods as a function of increasing input size. **(A, G)** Overall Elapsed Time: Total elapsed CPU time to complete processing. **(B, H)** Average Memory Usage: Mean CPU memory consumption (in MB) during execution, reflecting the typical memory footprint. **(C, I)** Peak Memory Usage: Maximum CPU memory recorded (in MB) at any point. **(D, J)** Peak GPU Memory: Maximum GPU memory recorded (in MB) at any point. **(E, K)** Memory Growth Rate: The average rate of memory increase during runtime, which indicates how the constant of memory usage increases with larger inputs. **(F, L)** CPU Utilization Profile: Percentage of time spent in native C execution (as opposed to interpreted Python code), reflecting the runtime that is being used by compiled, low-level routines.

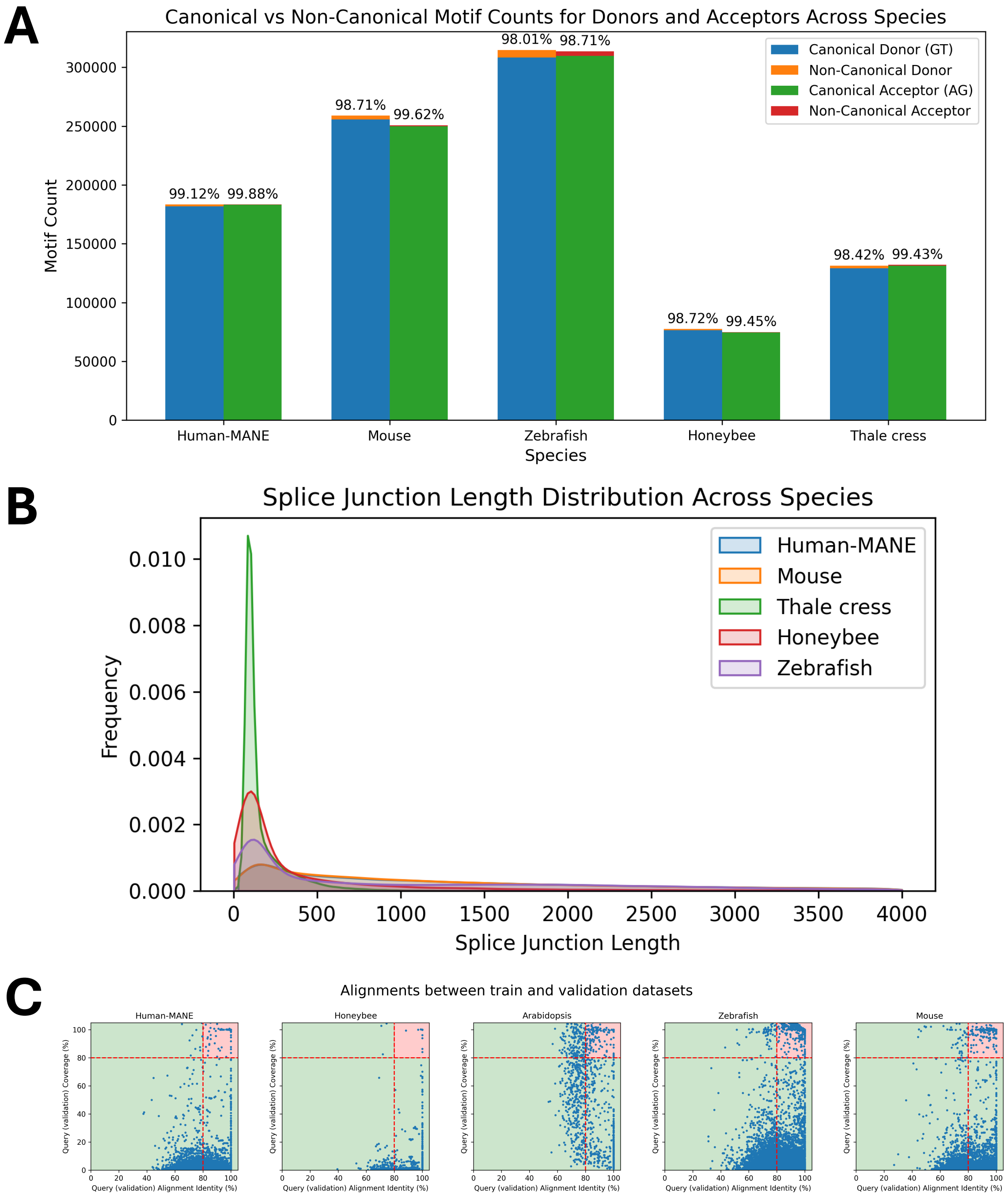


**Figure S7.** Splice site motif count and intron length distributions across five species. **(A)** Canonical vs. Non-Canonical Donor and Acceptor Splice Sites. Bar plots depict the total number of donor (blue, orange) and acceptor (green, red) sites across Human (MANE), Mouse, Zebrafish, Honeybee, and *Arabidopsis* genomes, subdivided into canonical (GT/AG) and non-canonical motifs. Percentages above each bar indicate the proportion of sites using canonical motifs in each species. **(B)** Splice Junction (Intron) Length Distributions. Kernel density curves illustrate the distribution of intron lengths in each genome (colored as in the legend). Differences in the breadth and peak of each distribution highlight notable cross-species variation in intron size. **(C)** Scatter plots of DNA sequence alignments between validation and training sets for Human-MANE, mouse, honeybee, zebrafish, and *Arabidopsis*. Each dot represents an alignment, with the x-axis showing alignment identity and the y-axis showing alignment coverage. Alignments exceeding 80% for both identity and coverage are highlighted in the red-shaded region and excluded from the test sets.

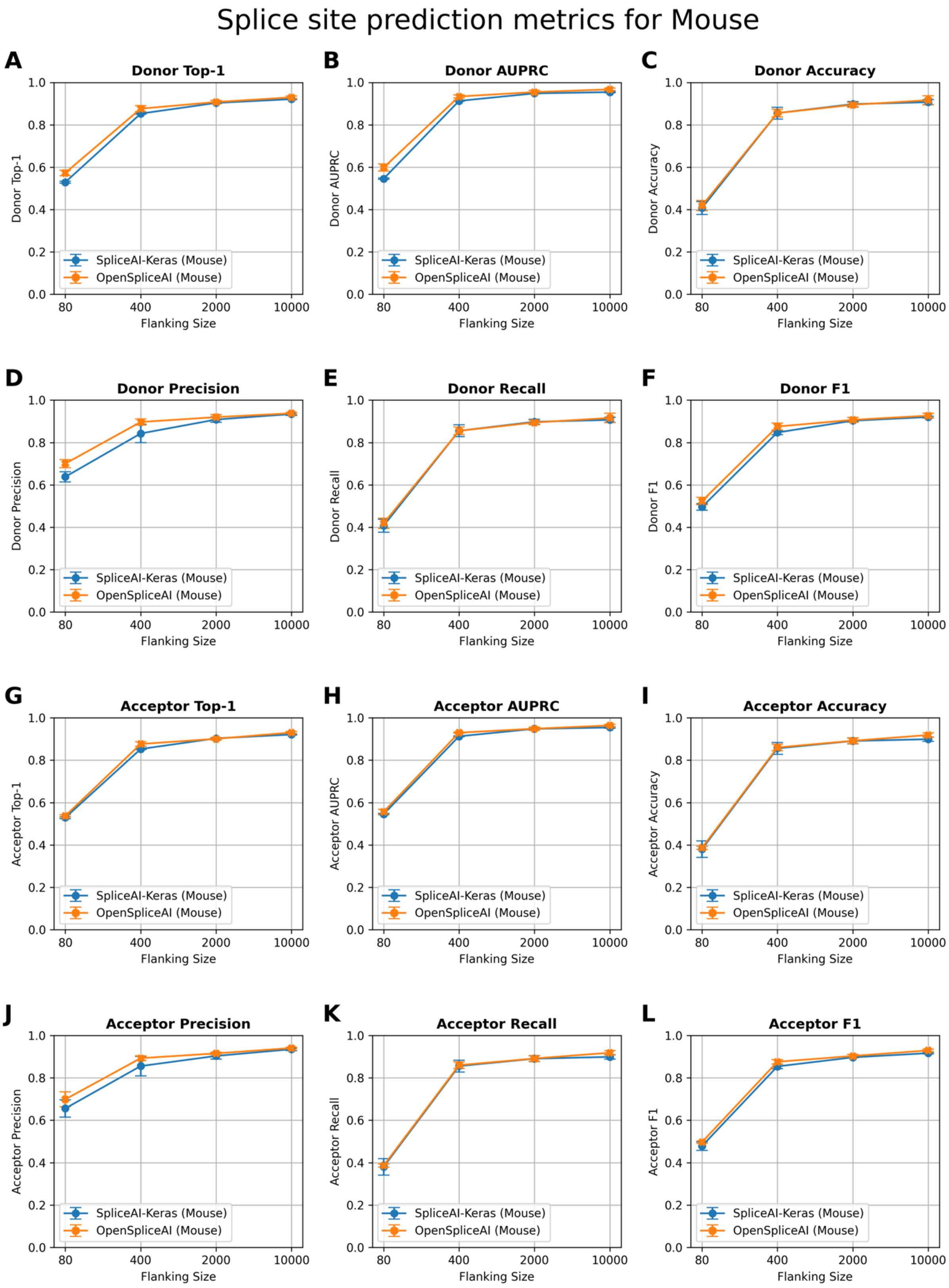


**Figure S8.** Splice site prediction metrics for the mouse (*Mus musculus*) across varying flanking sequence lengths. Plots compare performance of SpliceAI-Keras (blue) and $\text{OSAI}_{\text{Mouse}}$ (orange) on donor (5′) and acceptor (3′) splice site predictions using 80, 400, 2000, and 10,000 nt of flanking context. $\text{OSAI}_{\text{Mouse}}$ was trained with RefSeq GRCm39 annotation(https://ftp.ncbi.nlm.nih.gov/genomes/all/GCF/000/001/635/GCF_000001635.27_GRCm39/GCF_000001635.27_GRCm39_genomic.gff.gz) and GRCm39 genome. **(A)** Donor Top-1: measures the percentage of times the model's most confident prediction exactly matches the true donor site label. **(B)** Donor AUPRC: Area Under the Precision–Recall Curve for donor site predictions. **(C)** Donor Accuracy: proportion of correct donor site calls among all predictions. **(D)** Donor Precision: the fraction of predicted donor sites that are correct. **(E)** Donor Recall: the fraction of true donor sites that are correctly predicted. **(F)** Donor F1: the harmonic mean of precision and recall for donor sites. **(G)** Acceptor Top-1: measures the percentage of times the model's most confident prediction exactly matches the true acceptor site label. **(H)** Acceptor AUPRC: Area Under the Precision–Recall Curve for acceptor site predictions. **(I)** Acceptor Accuracy: proportion of correct acceptor site calls among all predictions. **(J)** Acceptor Precision: the fraction of predicted acceptor sites that are correct. **(K)** Acceptor Recall: the fraction of true acceptor sites that are correctly predicted. **(L)** Acceptor F1: the harmonic mean of precision and recall for acceptor sites. Each panel illustrates that increasing the flanking sequence length generally enhances model performance, with both SpliceAI-Keras and OpenSpliceAI achieving high accuracy and F1 scores at 10,000 nt. Each data point represents the mean across five independently trained models, with error bars indicating the standard deviation.

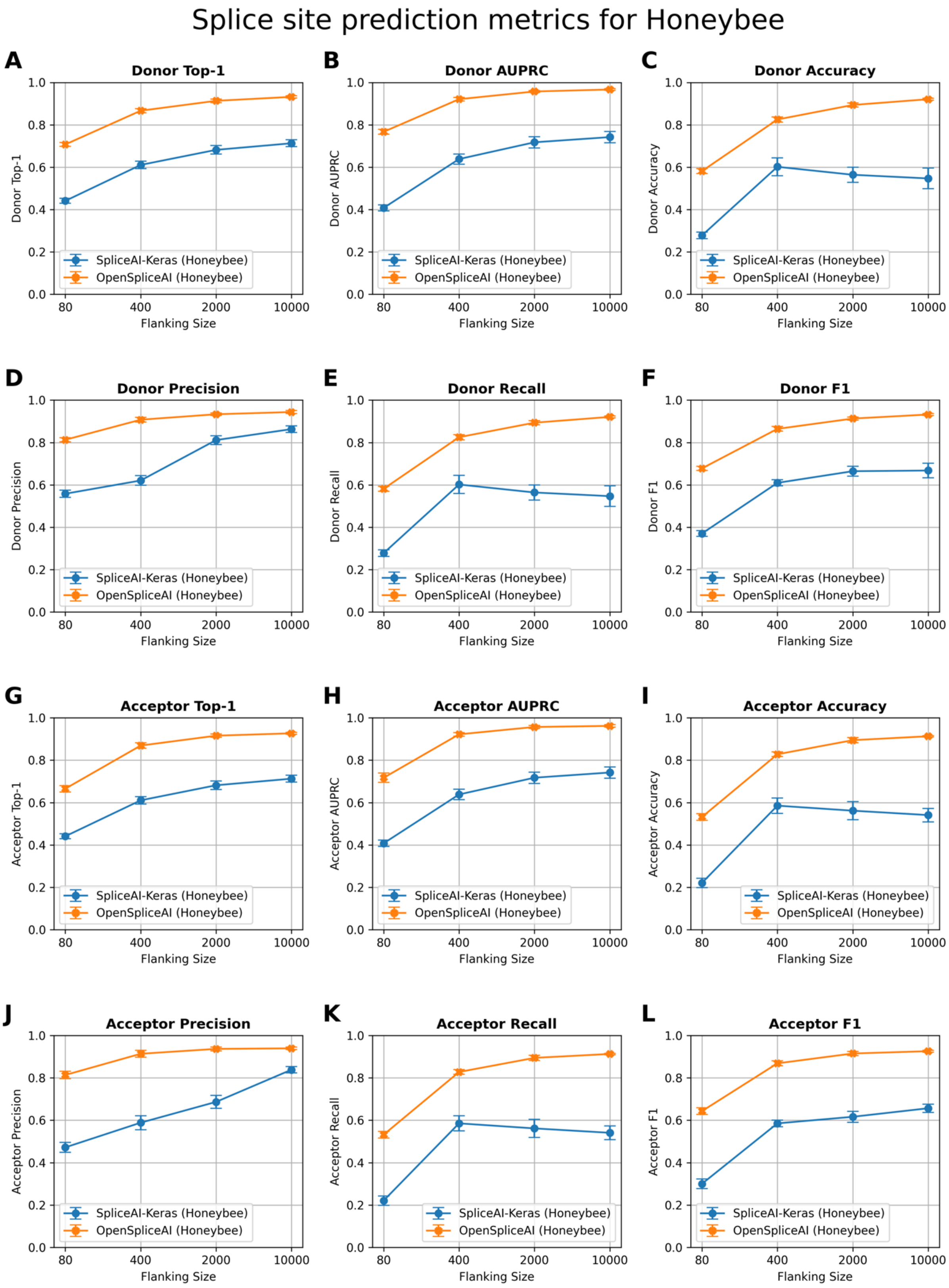


**Figure S9.** Splice site prediction metrics for the honeybee (*Apis mellifera*) across varying flanking sequence lengths. Plots compare performance of SpliceAI-Keras (blue) and $OSAI_{Honeybee}$ (orange) on donor (5′) and acceptor (3′) splice site predictions using 80, 400, 2000, and 10,000 nt of flanking context. $OSAI_{Honeybee}$ was trained with RefSeq Amel_HAv3.1 annotation (https://ftp.ncbi.nlm.nih.gov/genomes/all/GCF/003/254/395/GCF_003254395.2_Amel_HAv3.1/GCF_003254395.2_Amel_HAv3.1_genomic.gff.gz) and Amel_HAv3.1 genome. **(A)** Donor Top-1: measures the percentage of times the model's most confident prediction exactly matches the true donor site label. **(B)** Donor AUPRC: Area Under the Precision–Recall Curve for donor site predictions. **(C)** Donor Accuracy: proportion of correct donor site calls among all predictions. **(D)** Donor Precision: the fraction of predicted donor sites that are correct. **(E)** Donor Recall: the fraction of true donor sites that are correctly predicted. **(F)** Donor F1: the harmonic mean of precision and recall for donor sites. **(G)** Acceptor Top-1: measures the percentage of times the model's most confident prediction exactly matches the true acceptor site label. **(H)** Acceptor AUPRC: Area Under the Precision–Recall Curve for acceptor site predictions. **(I)** Acceptor Accuracy: proportion of correct acceptor site calls among all predictions. **(J)** Acceptor Precision: the fraction of predicted acceptor sites that are correct. **(K)** Acceptor Recall: the fraction of true acceptor sites that are correctly predicted. **(L)** Acceptor F1: the harmonic mean of precision and recall for acceptor sites. Each panel illustrates that increasing the flanking sequence length generally enhances model performance, with both SpliceAI-Keras and OpenSpliceAI achieving high accuracy and F1 scores at 10,000 nt. Each data point represents the mean across five independently trained models, with error bars indicating the standard deviation.

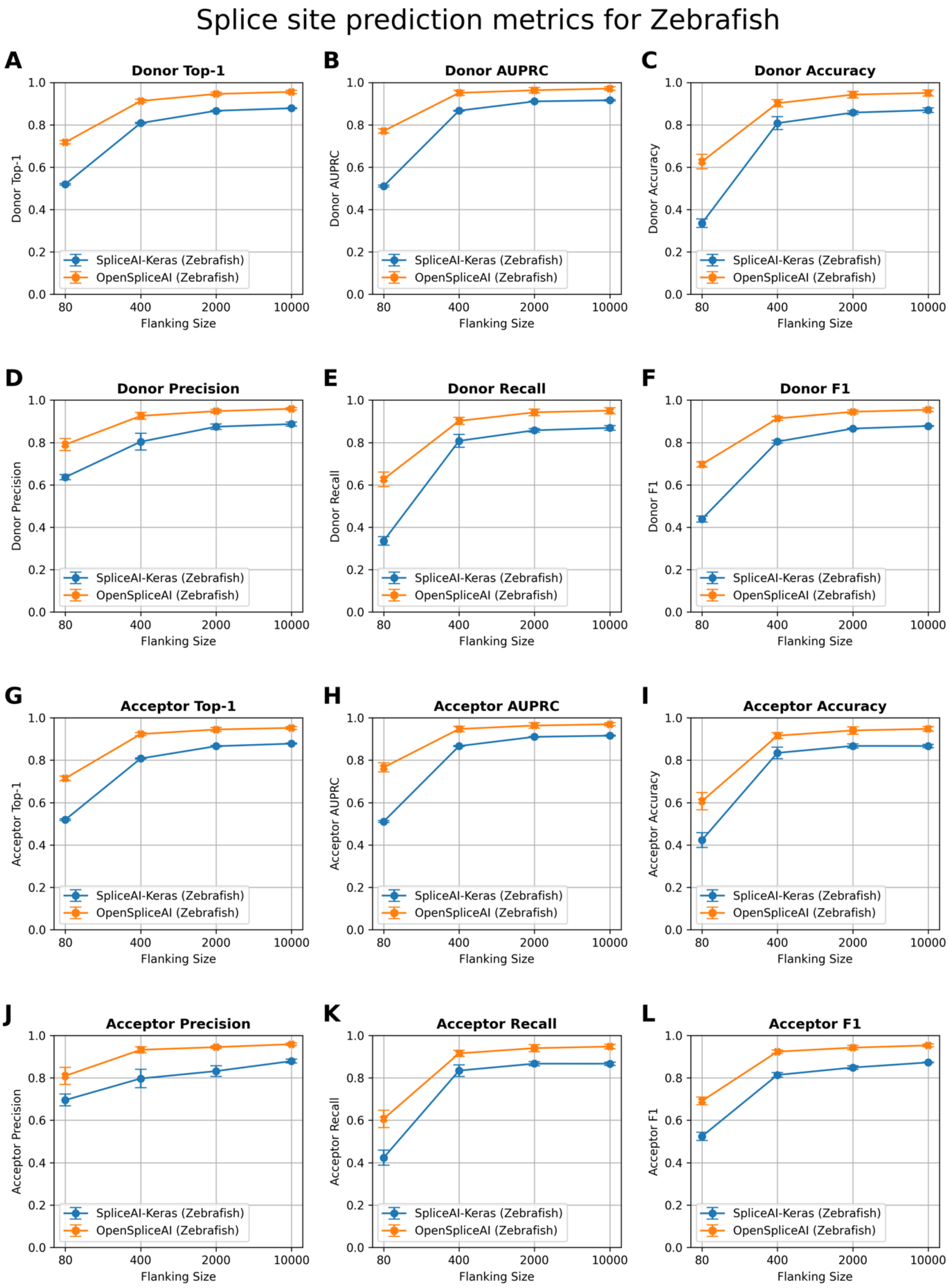


**Figure S10.** Splice site prediction metrics for the zebrafish (*Danio rerio*) across varying flanking sequence lengths. Plots compare performance of SpliceAI-Keras (blue) and $OSAI_{Zebrafish}$ (orange) on donor (5′) and acceptor (3′) splice site predictions using 80, 400, 2000, and 10,000 nt of flanking context. $OSAI_{Zebrafish}$ was trained with RefSeq GRCz11 annotation (https://ftp.ncbi.nlm.nih.gov/genomes/all/GCF/000/002/035/GCF_000002035.6_GRCz11/GCF_000002035.6_GRCz11_genomic.gff.gz) and GRCz11 genome. **(A)** Donor Top-1: measures the percentage of times the model's most confident prediction exactly matches the true donor site label. **(B)** Donor AUPRC: Area Under the Precision–Recall Curve for donor site predictions. **(C)** Donor Accuracy: proportion of correct donor site calls among all predictions. **(D)** Donor Precision: the fraction of predicted donor sites that are correct. **(E)** Donor Recall: the fraction of true donor sites that are correctly predicted. **(F)** Donor F1: the harmonic mean of precision and recall for donor sites. **(G)** Acceptor Top-1: measures the percentage of times the model's most confident prediction exactly matches the true acceptor site label. **(H)** Acceptor AUPRC: Area Under the Precision–Recall Curve for acceptor site predictions. **(I)** Acceptor Accuracy: proportion of correct acceptor site calls among all predictions. **(J)** Acceptor Precision: the fraction of predicted acceptor sites that are correct. **(K)** Acceptor Recall: the fraction of true acceptor sites that are correctly predicted. **(L)** Acceptor F1: the harmonic mean of precision and recall for acceptor sites. Each panel illustrates that increasing the flanking sequence length generally enhances model performance, with both SpliceAI-Keras and OpenSpliceAI achieving high accuracy and F1 scores at 10,000 nt. Each data point represents the mean across five independently trained models, with error bars indicating the standard deviation.

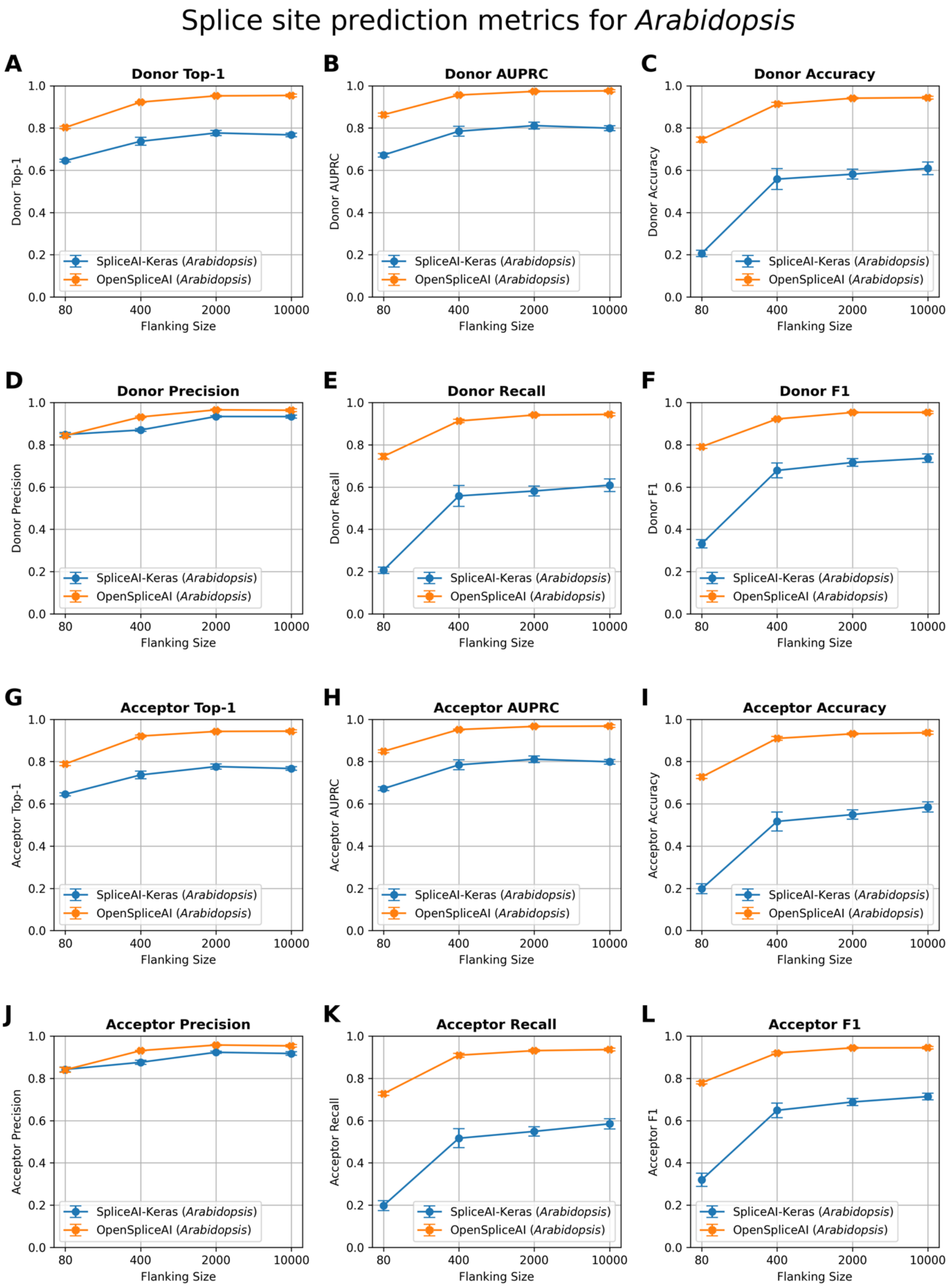


**Figure S11.** Splice site prediction metrics for the *Arabidopsis thaliana* across varying flanking sequence lengths. Plots compare performance of SpliceAI-Keras (blue) and $OSAI_{Arabidopsis}$ (orange) on donor (5′) and acceptor (3′) splice site predictions using 80, 400, 2000, and 10,000 nt of flanking context. $OSAI_{Arabidopsis}$

was trained with RefSeq TAIR10.1 annotation (https://ftp.ncbi.nlm.nih.gov/genomes/all/GCF/000/001/735/GCF_000001735.4_TAIR10.1/GCF_000001735.4_TAIR10.1_genomic.gff.gz) and TAIR10.1 genome. **(A)** Donor Top-1: measures the percentage of times the model's most confident prediction exactly matches the true donor site label. **(B)** Donor AUPRC: Area Under the Precision–Recall Curve for donor site predictions. **(C)** Donor Accuracy: proportion of correct donor site calls among all predictions. **(D)** Donor Precision: the fraction of predicted donor sites that are correct. **(E)** Donor Recall: the fraction of true donor sites that are correctly predicted. **(F)** Donor F1: the harmonic mean of precision and recall for donor sites. **(G)** Acceptor Top-1: measures the percentage of times the model's most confident prediction exactly matches the true acceptor site label. **(H)** Acceptor AUPRC: Area Under the Precision–Recall Curve for acceptor site predictions. **(I)** Acceptor Accuracy: proportion of correct acceptor site calls among all predictions. **(J)** Acceptor Precision: the fraction of predicted acceptor sites that are correct. **(K)** Acceptor Recall: the fraction of true acceptor sites that are correctly predicted. **(L)** Acceptor F1: the harmonic mean of precision and recall for acceptor sites. Each panel illustrates that increasing the flanking sequence length generally enhances model performance, with both SpliceAI-Keras and OpenSpliceAI achieving high accuracy and F1 scores at 10,000 nt. Each data point represents the mean across five independently trained models, with error bars indicating the standard deviation.

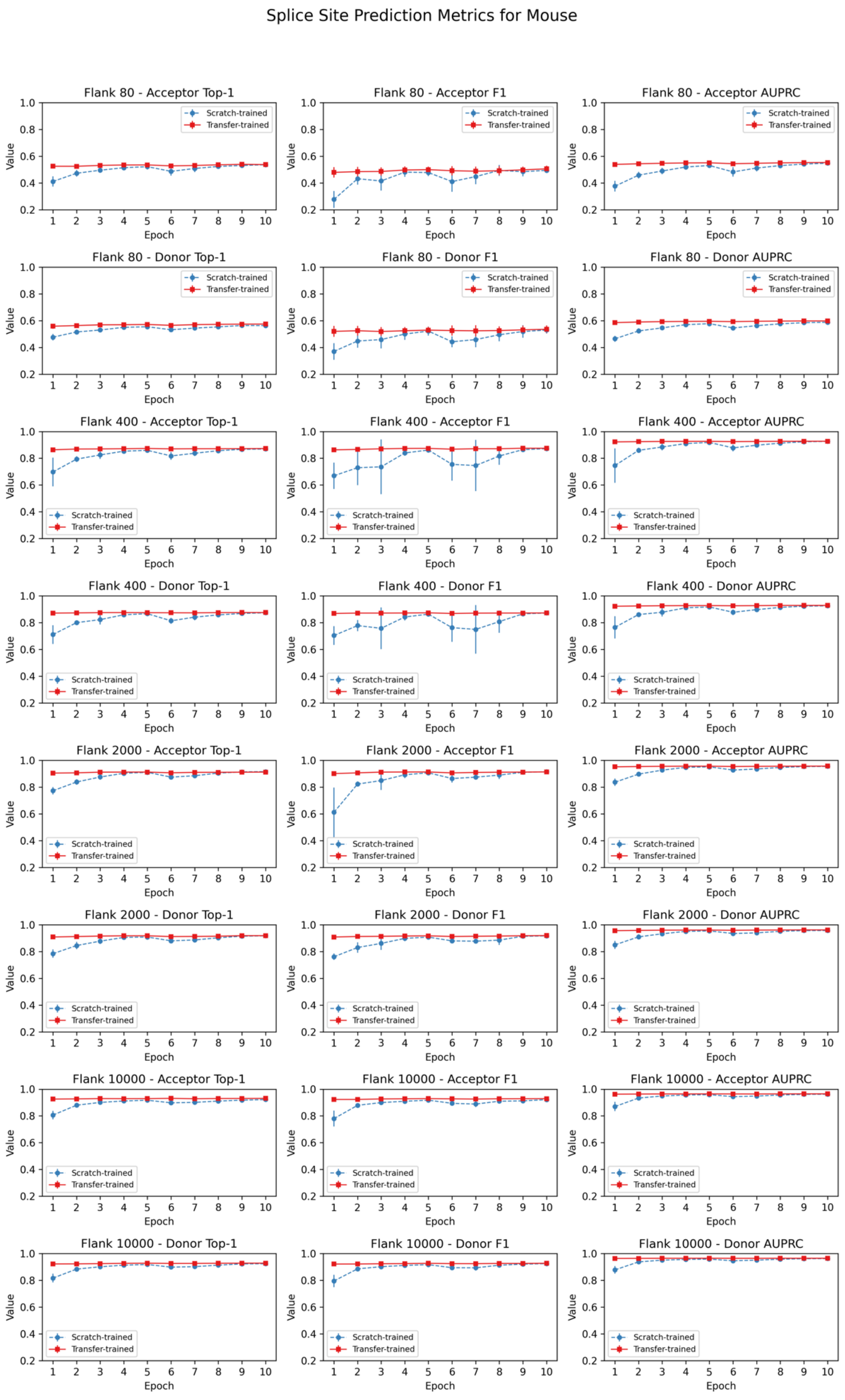
Splice Site Prediction Metrics for Mouse
Flank 80 - Acceptor Top-1
Flank 80 - Acceptor F1
Flank 80 - Acceptor AUPRC
Flank 80 - Donor Top-1
Flank 80 - Donor F1
Flank 80 - Donor AUPRC
Flank 400 - Acceptor Top-1
Flank 400 - Acceptor F1
Flank 400 - Acceptor AUPRC
Flank 400 - Donor Top-1
Flank 400 - Donor F1
Flank 400 - Donor AUPRC
Flank 2000 - Acceptor Top-1
Flank 2000 - Acceptor F1
Flank 2000 - Acceptor AUPRC
Flank 2000 - Donor Top-1
Flank 2000 - Donor F1
Flank 2000 - Donor AUPRC
Flank 10000 - Acceptor Top-1
Flank 10000 - Acceptor F1
Flank 10000 - Acceptor AUPRC
Flank 10000 - Donor Top-1
Flank 10000 - Donor F1
Flank 10000 - Donor AUPRC
Scratch-trained
Transfer-trained
Value
Epoch

**Figure S12.** Transfer learning from $OSAI_{MANE}$ was leveraged to evaluate performance metrics for splice site prediction in mouse (*Mus musculus*) across four flanking sequence lengths (80, 400, 2,000, and 10,000 nucleotides) and two splice site types (acceptor and donor). Each row corresponds to a unique flanking length–splice site combination, and each column depicts a distinct evaluation metric (Top-1 accuracy, F1, and AUPRC). Blue and red curves represent the mean performance (± standard deviation) of five models trained from scratch and five models fine-tuned from $OSAI_{MANE}$, respectively. The x-axis indicates the training epoch, and the y-axis denotes the corresponding metric value. Subplot titles specify the flanking length, splice site type, and metric. Overall, fine-tuned models converge more rapidly, reaching stable performance as early as the first epoch, whereas models trained from scratch require additional epochs to stabilize.

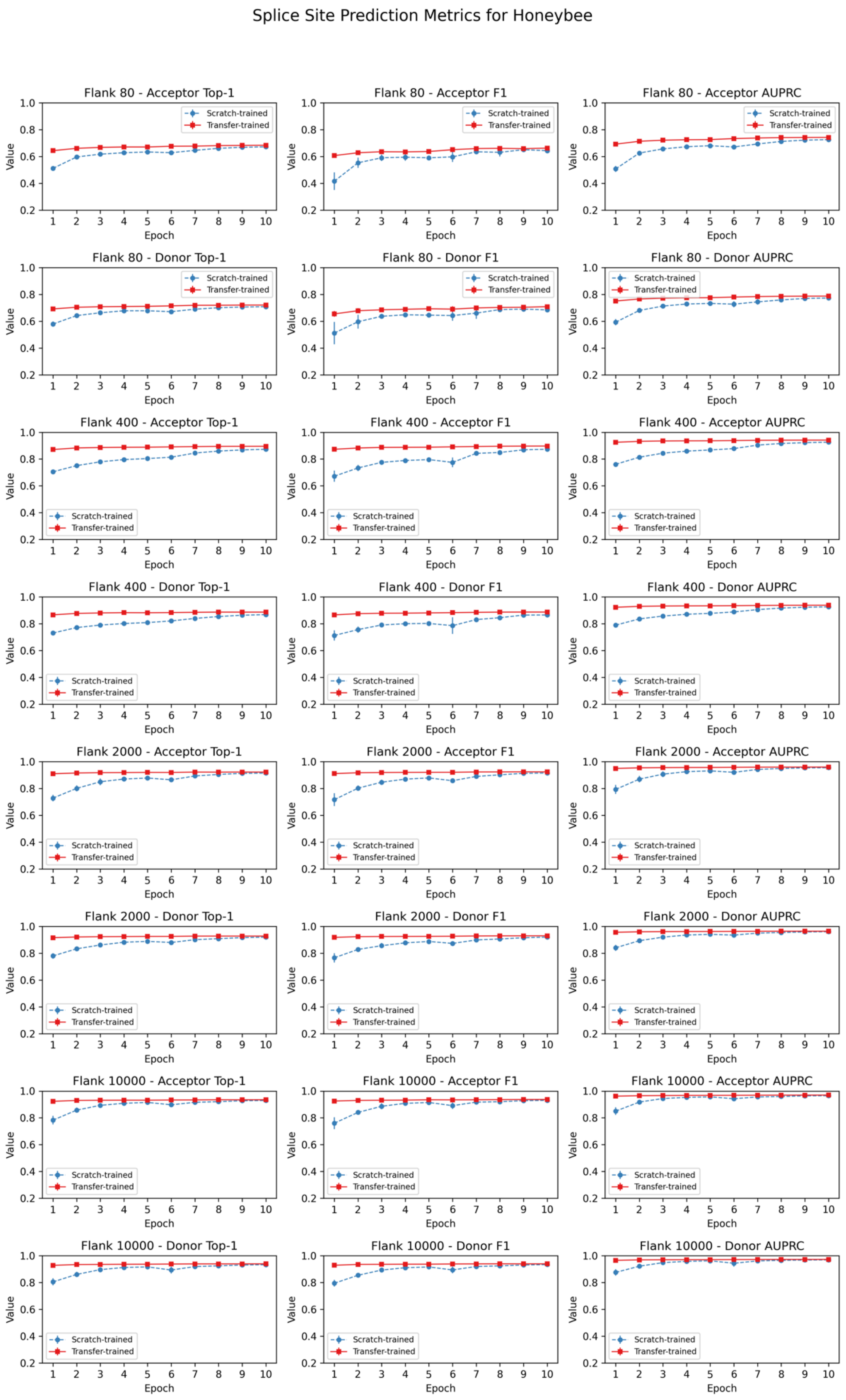
Splice Site Prediction Metrics for Honeybee
Flank 80 - Acceptor Top-1
Flank 80 - Acceptor F1
Flank 80 - Acceptor AUPRC
Flank 80 - Donor Top-1
Flank 80 - Donor F1
Flank 80 - Donor AUPRC
Flank 400 - Acceptor Top-1
Flank 400 - Acceptor F1
Flank 400 - Acceptor AUPRC
Flank 400 - Donor Top-1
Flank 400 - Donor F1
Flank 400 - Donor AUPRC
Flank 2000 - Acceptor Top-1
Flank 2000 - Acceptor F1
Flank 2000 - Acceptor AUPRC
Flank 2000 - Donor Top-1
Flank 2000 - Donor F1
Flank 2000 - Donor AUPRC
Flank 10000 - Acceptor Top-1
Flank 10000 - Acceptor F1
Flank 10000 - Acceptor AUPRC
Flank 10000 - Donor Top-1
Flank 10000 - Donor F1
Flank 10000 - Donor AUPRC
Scratch-trained
Transfer-trained
Value
Epoch

**Figure S13.** Transfer learning from $OSAI_{MANE}$ was leveraged to evaluate performance metrics for splice site prediction in honeybee (*Apis mellifera*) across four flanking sequence lengths (80, 400, 2,000, and 10,000 nucleotides) and two splice site types (acceptor and donor). Each row corresponds to a unique flanking length–splice site combination, and each column depicts a distinct evaluation metric (Top-1 accuracy, F1, and AUPRC). Blue and red curves represent the mean performance ($\pm$ standard deviation) of five models trained from scratch and five models fine-tuned from $OSAI_{MANE}$, respectively. The x-axis indicates the training epoch, and the y-axis denotes the corresponding metric value. Subplot titles specify the flanking length, splice site type, and metric. Overall, fine-tuned models converge more rapidly, reaching stable performance as early as the first epoch, whereas models trained from scratch require additional epochs to stabilize.

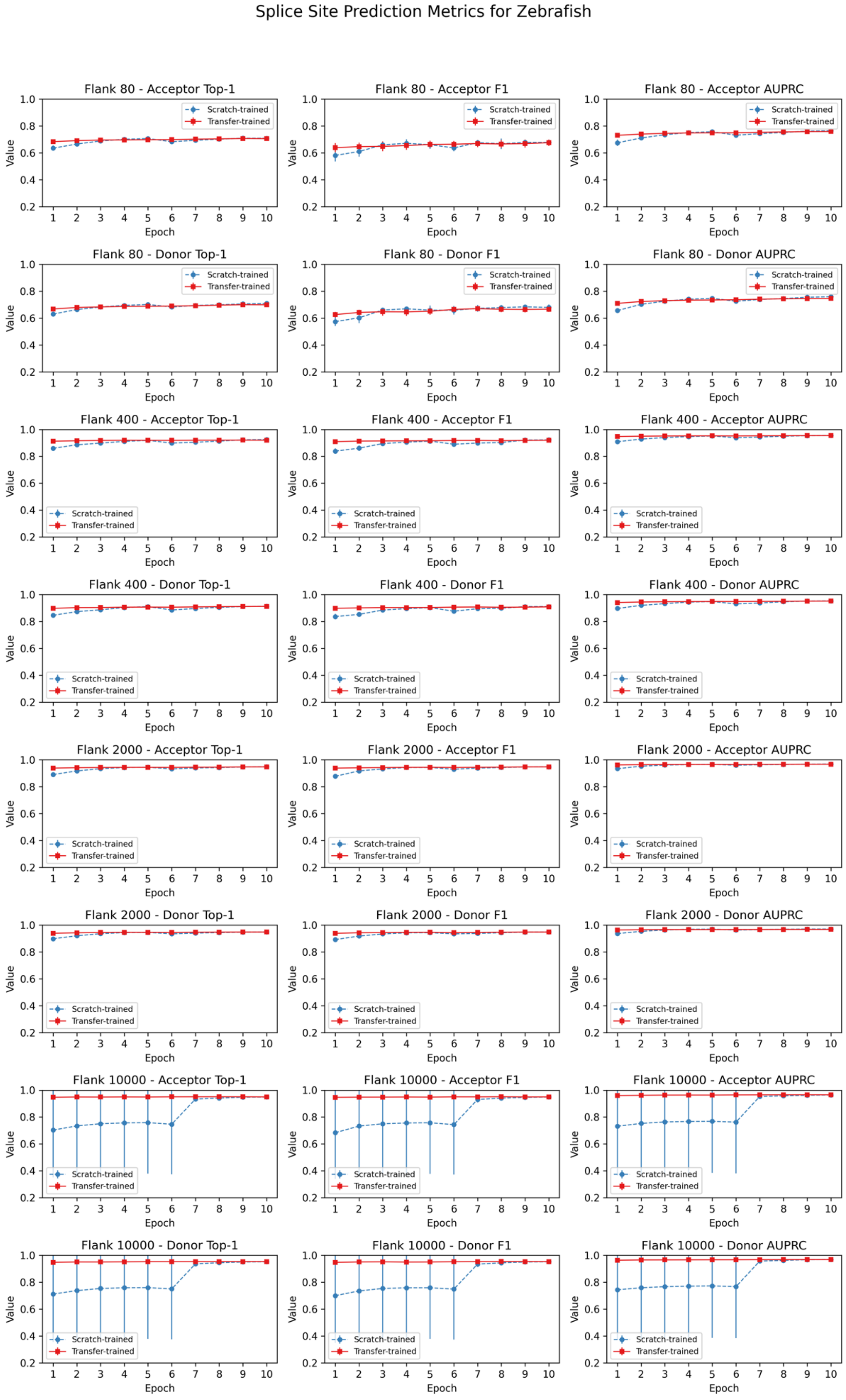
Splice Site Prediction Metrics for Zebrafish
Flank 80 - Acceptor Top-1
Flank 80 - Acceptor F1
Flank 80 - Acceptor AUPRC
Flank 80 - Donor Top-1
Flank 80 - Donor F1
Flank 80 - Donor AUPRC
Flank 400 - Acceptor Top-1
Flank 400 - Acceptor F1
Flank 400 - Acceptor AUPRC
Flank 400 - Donor Top-1
Flank 400 - Donor F1
Flank 400 - Donor AUPRC
Flank 2000 - Acceptor Top-1
Flank 2000 - Acceptor F1
Flank 2000 - Acceptor AUPRC
Flank 2000 - Donor Top-1
Flank 2000 - Donor F1
Flank 2000 - Donor AUPRC
Flank 10000 - Acceptor Top-1
Flank 10000 - Acceptor F1
Flank 10000 - Acceptor AUPRC
Flank 10000 - Donor Top-1
Flank 10000 - Donor F1
Flank 10000 - Donor AUPRC
Scratch-trained
Transfer-trained
Value
Epoch

**Figure S14.** Transfer learning from $OSAI_{MANE}$ was leveraged to evaluate performance metrics for splice site prediction in zebrafish (*Danio rerio*) across four flanking sequence lengths (80, 400, 2,000, and 10,000 nucleotides) and two splice site types (acceptor and donor). Each row corresponds to a unique flanking length–splice site combination, and each column depicts a distinct evaluation metric (Top-1 accuracy, F1, and AUPRC). Blue and red curves represent the mean performance ($\pm$ standard deviation) of five models trained from scratch and five models fine-tuned from $OSAI_{MANE}$, respectively. The x-axis indicates the training epoch, and the y-axis denotes the corresponding metric value. Subplot titles specify the flanking length, splice site type, and metric. Overall, fine-tuned models converge more rapidly, reaching stable performance as early as the first epoch, whereas models trained from scratch require additional epochs to stabilize.

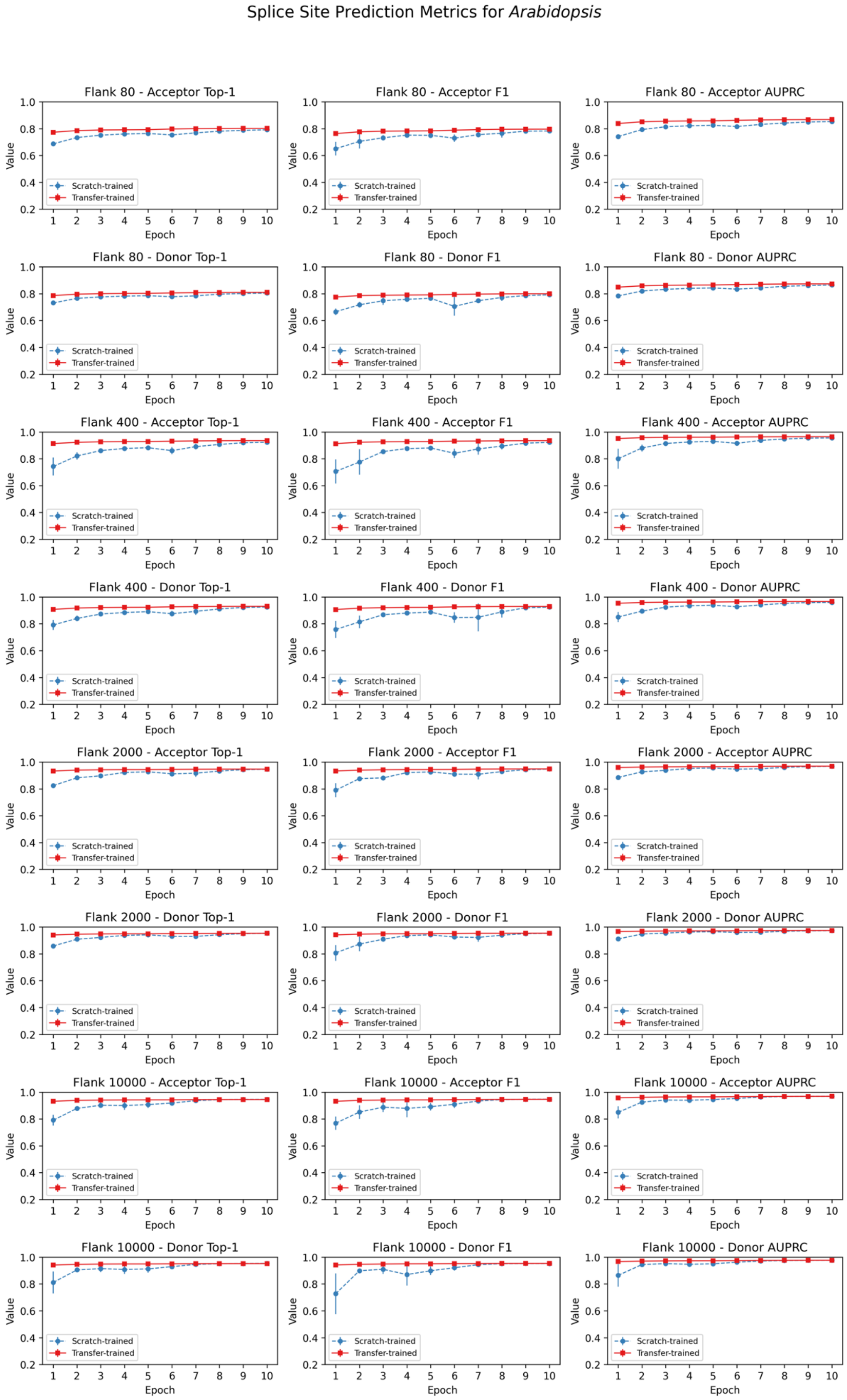
Splice Site Prediction Metrics for *Arabidopsis*
Flank 80 - Acceptor Top-1
Flank 80 - Acceptor F1
Flank 80 - Acceptor AUPRC
Flank 80 - Donor Top-1
Flank 80 - Donor F1
Flank 80 - Donor AUPRC
Flank 400 - Acceptor Top-1
Flank 400 - Acceptor F1
Flank 400 - Acceptor AUPRC
Flank 400 - Donor Top-1
Flank 400 - Donor F1
Flank 400 - Donor AUPRC
Flank 2000 - Acceptor Top-1
Flank 2000 - Acceptor F1
Flank 2000 - Acceptor AUPRC
Flank 2000 - Donor Top-1
Flank 2000 - Donor F1
Flank 2000 - Donor AUPRC
Flank 10000 - Acceptor Top-1
Flank 10000 - Acceptor F1
Flank 10000 - Acceptor AUPRC
Flank 10000 - Donor Top-1
Flank 10000 - Donor F1
Flank 10000 - Donor AUPRC
Scratch-trained
Transfer-trained
Value
Epoch

**Figure S15.** Transfer learning from $OSAI_{MANE}$ was leveraged to evaluate performance metrics for splice site prediction in *Arabidopsis thaliana* across four flanking sequence lengths (80, 400, 2,000, and 10,000 nucleotides) and two splice site types (acceptor and donor). Each row corresponds to a unique flanking length–splice site combination, and each column depicts a distinct evaluation metric (Top-1 accuracy, F1, and AUPRC). Blue and red curves represent the mean performance (± standard deviation) of five models trained from scratch and five models fine-tuned from $OSAI_{MANE}$, respectively. The x-axis indicates the training epoch, and the y-axis denotes the corresponding metric value. Subplot titles specify the flanking length, splice site type, and metric. Overall, fine-tuned models converge more rapidly, reaching stable performance as early as the first epoch, whereas models trained from scratch require additional epochs to stabilize.

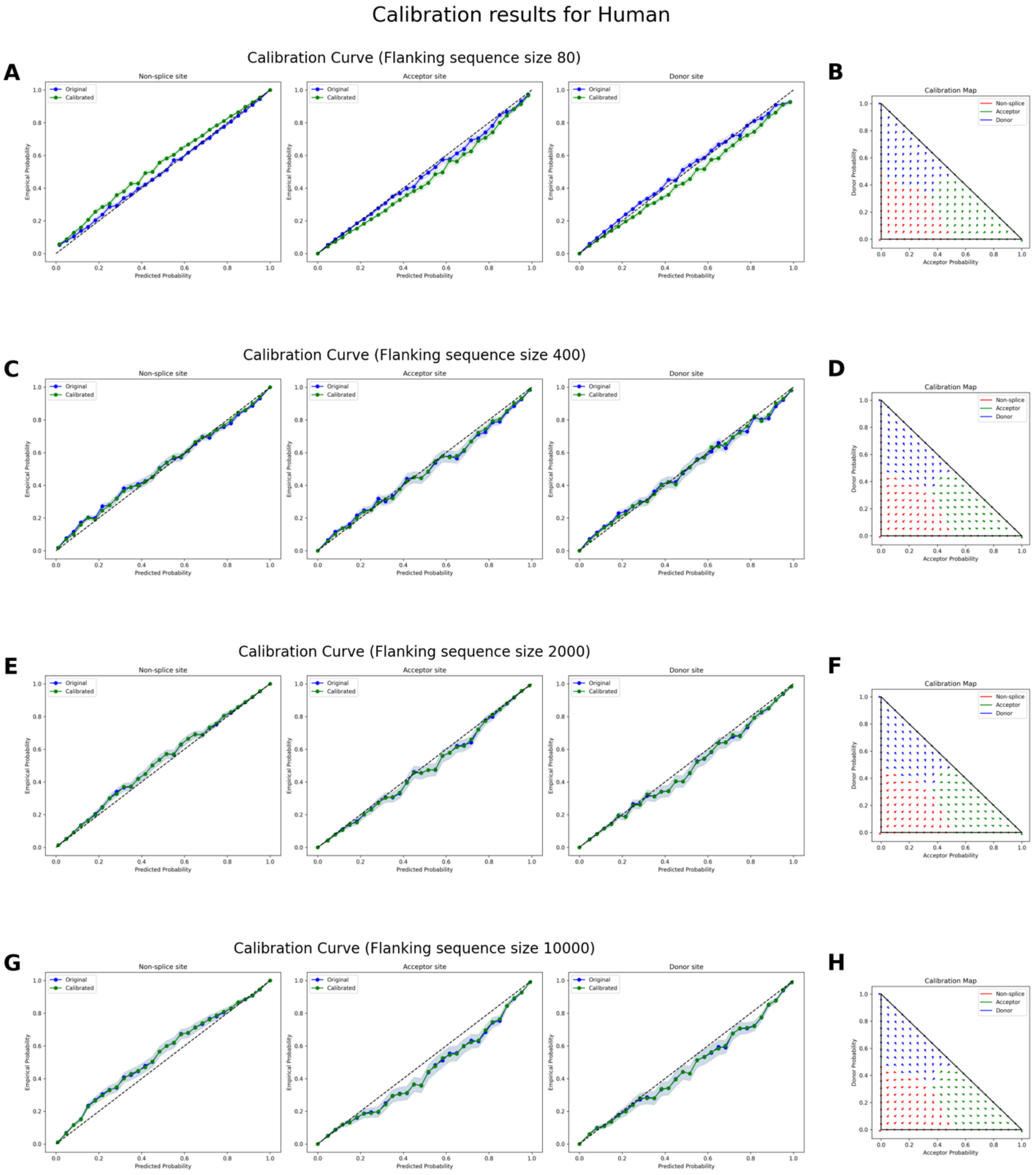


**Figure S16.** Calibration results for human MANE splice site classification at four flanking sequence sizes. **(A, C, E, G)** Reliability (calibration) curves for flanking sequence sizes of 80, 400, 2000, and 10,000 nucleotides, respectively. Each plot compares predicted probabilities (x-axis) to empirical probabilities (y-axis) for non-splice sites (left), acceptor sites (middle), and donor sites (right). The blue curves depict the reliability of the original OSAI$_{MANE}$ models, while the green curves show reliability after calibration. Shaded regions represent confidence intervals. The diagonal black line indicates perfect calibration, where predicted probabilities match observed frequencies exactly. **(B, D, F, H)** Temperature scaling maps for each

corresponding flanking sequence size, illustrating how raw predicted probabilities for acceptor (x-axis) and donor (y-axis) sites are transformed after calibration. Arrows indicate the shift from pre- to post-calibration states in two-dimensional probability space.

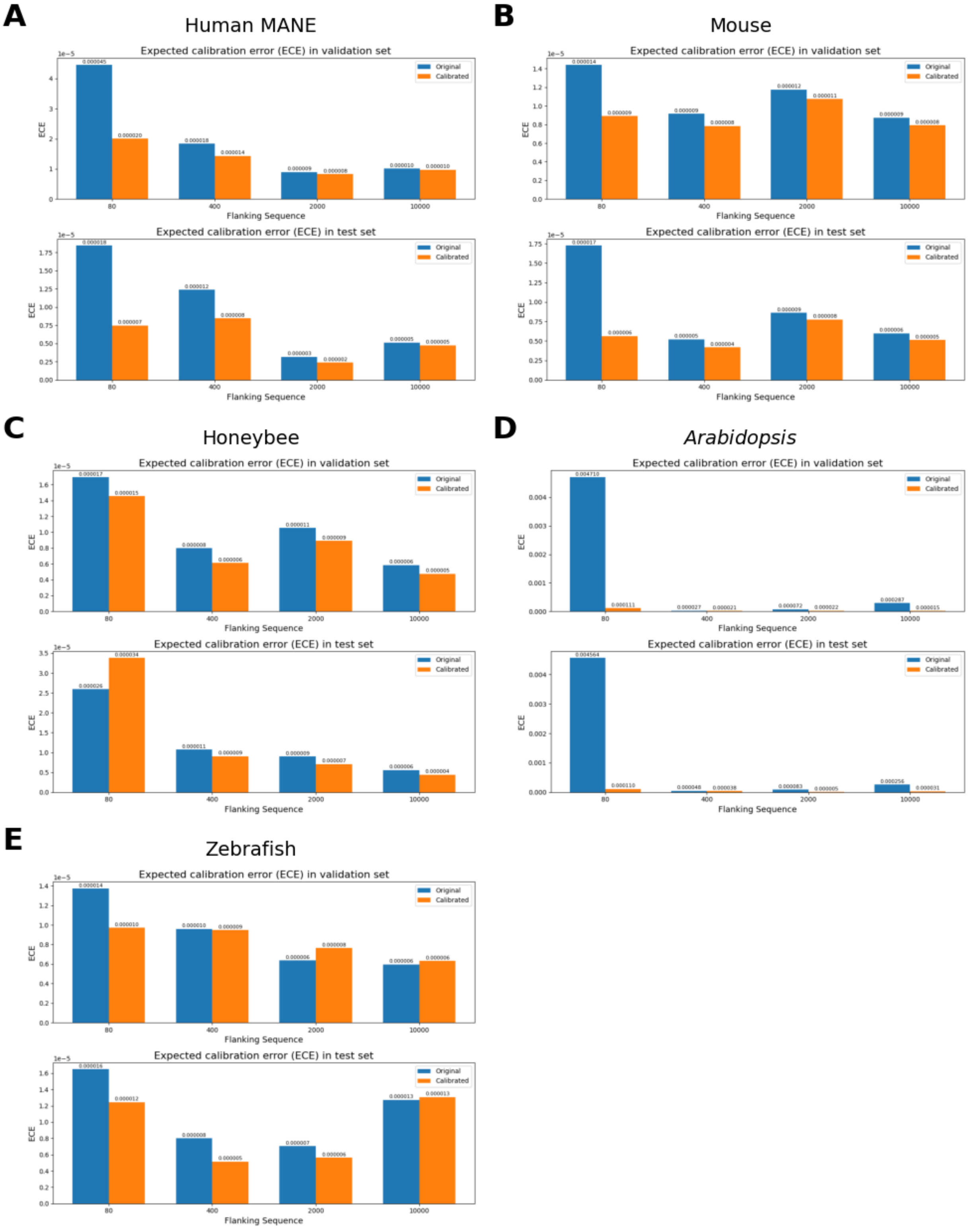


**Figure S17.** Expected Calibration Error (ECE) on the validation (top) and test (bottom) sets. Blue bars indicate model performance before calibration, and orange bars indicate performance after calibration. Results are shown for (A) Human-MANE, (B) mouse, (C) honeybee, (D) *Arabidopsis*, and (E) zebrafish.

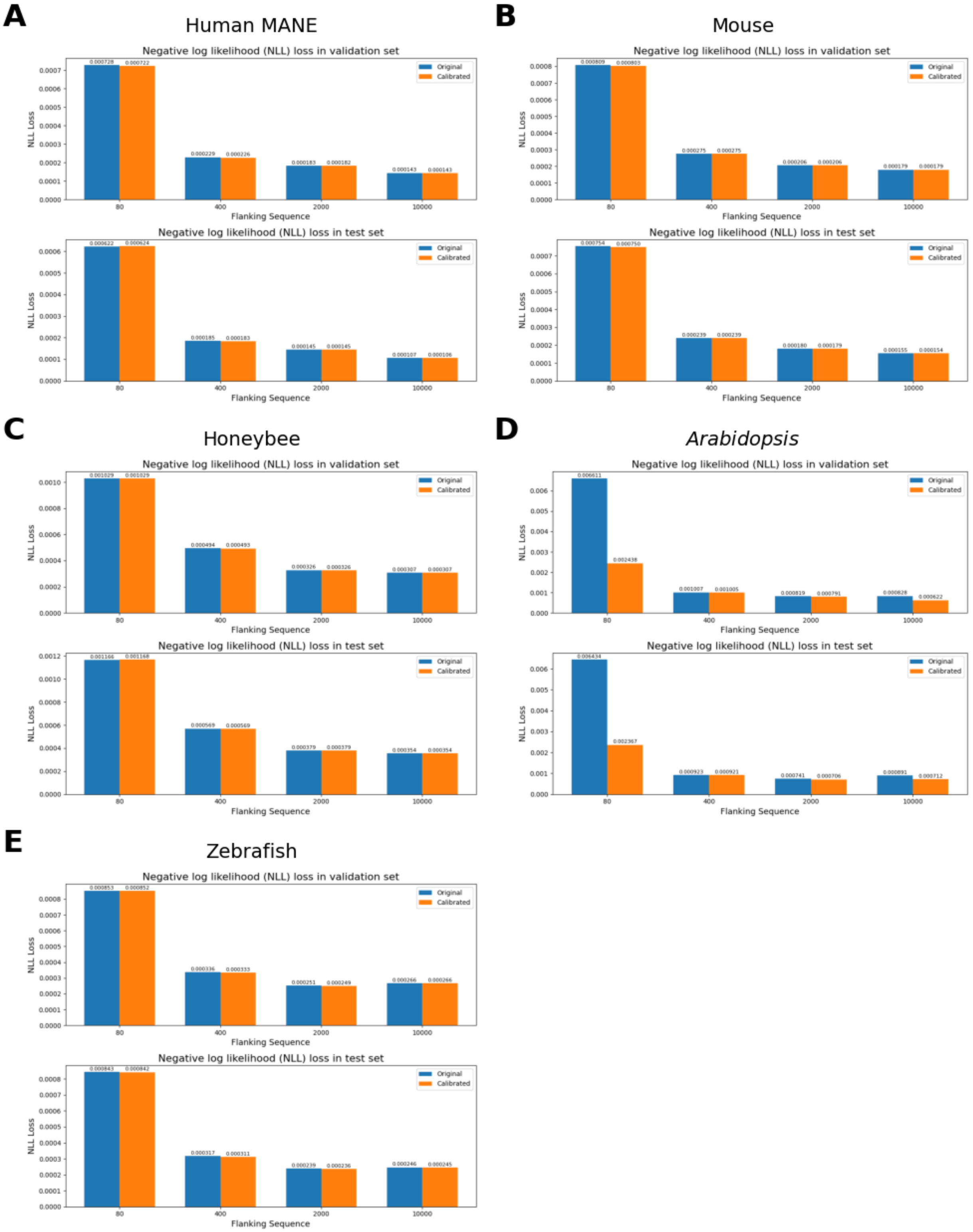


**Figure S18.** Negative log likelihood (NLL) loss on the validation (top) and test (bottom) sets. Blue bars indicate model performance before calibration, and orange bars indicate performance after calibration. Results are shown for (A) Human-MANE, (B) mouse, (C) honeybee, (D) *Arabidopsis*, and (E) zebrafish.

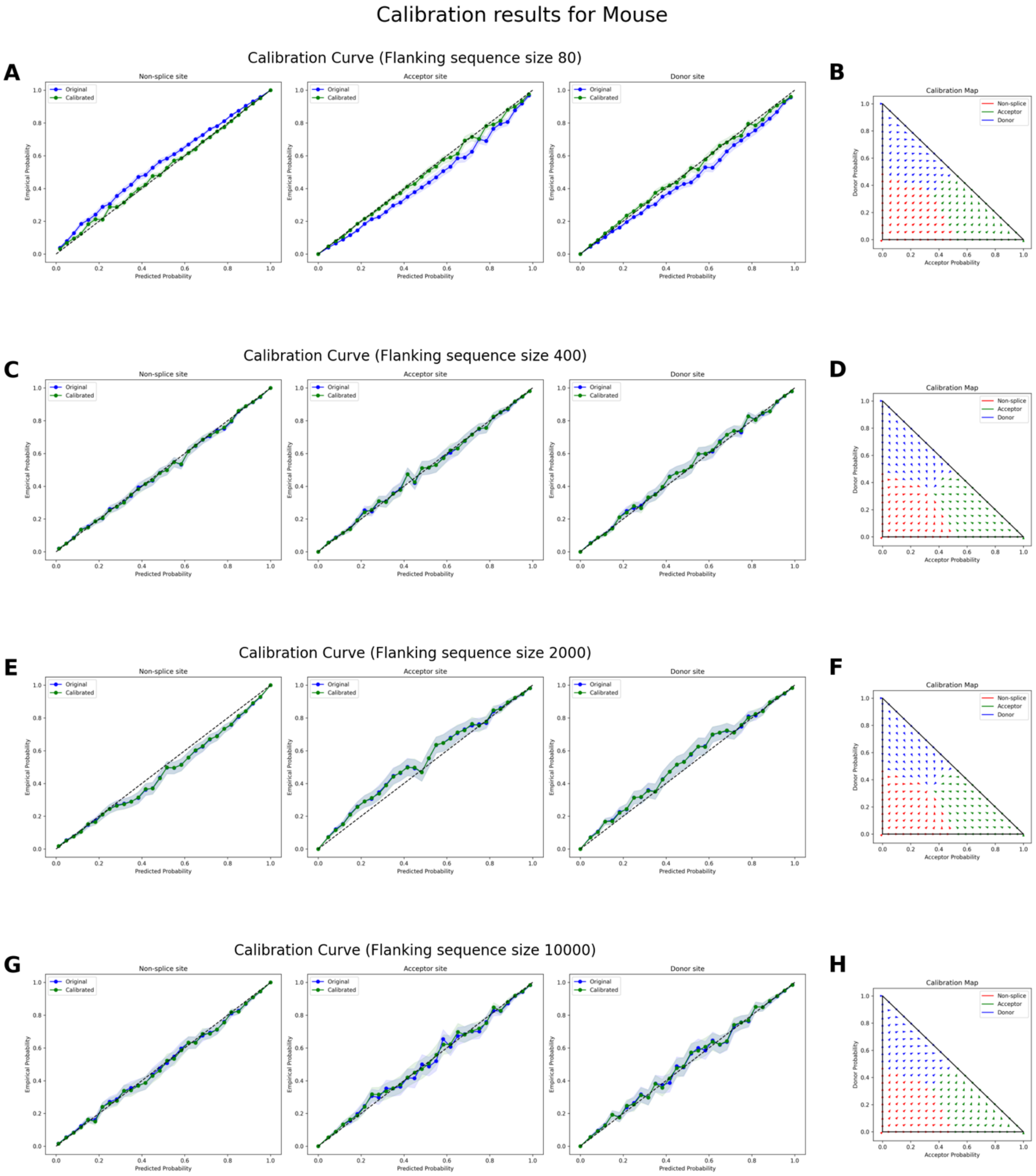


**Figure S19.** Calibration results for house mouse (*Mus musculus*) splice site classification at four flanking sequence sizes. **(A, C, E, G)** Reliability (calibration) curves for flanking sequence sizes of 80, 400, 2000, and 10,000 nucleotides, respectively. Each plot compares predicted probabilities (x-axis) to empirical probabilities (y-axis) for non-splice sites (left), acceptor sites (middle), and donor sites (right). The blue curves depict the reliability of the original $\text{OSAI}_{\text{Mouse}}$ models, while the green curves show reliability after calibration. Shaded regions represent confidence intervals. The diagonal black line indicates perfect calibration, where predicted probabilities match observed frequencies exactly. **(B, D, F, H)** Temperature scaling maps for each corresponding flanking sequence size, illustrating how raw predicted probabilities for acceptor (x-axis) and donor (y-axis) sites are transformed after calibration. Arrows indicate the shift from pre- to post-calibration states in two-dimensional probability space.

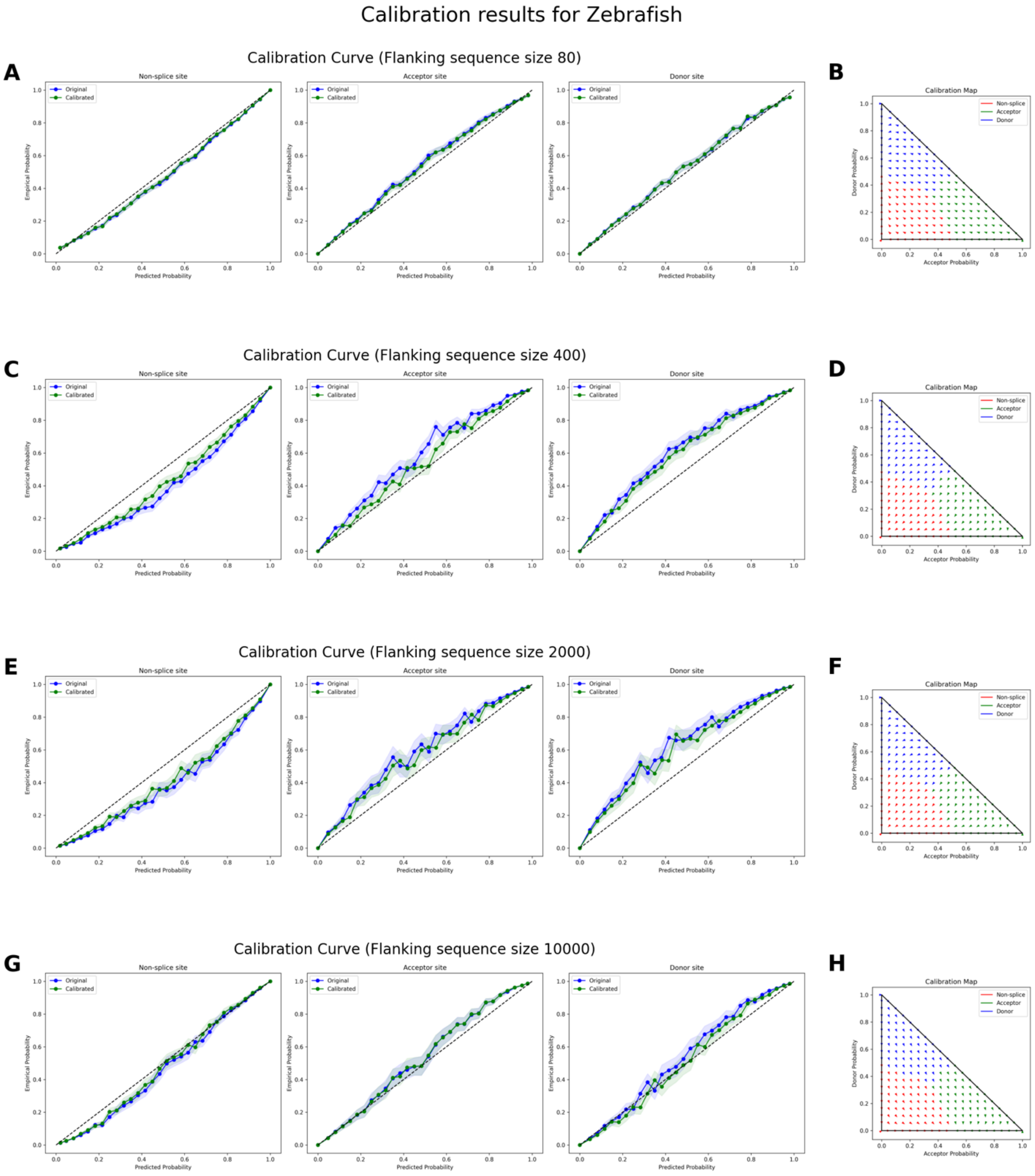


**Figure S20.** Calibration results for zebrafish (*Danio rerio*) splice site classification at four flanking sequence sizes. **(A, C, E, G)** Reliability (calibration) curves for flanking sequence sizes of 80, 400, 2000, and 10,000 nucleotides, respectively. Each plot compares predicted probabilities (x-axis) to empirical probabilities (y-axis) for non-splice sites (left), acceptor sites (middle), and donor sites (right). The blue curves depict the reliability of the original $OSAI_{Zebrafish}$ models, while the green curves show reliability after calibration. Shaded regions represent confidence intervals. The diagonal black line indicates perfect calibration, where predicted probabilities match observed frequencies exactly. **(B, D, F, H)** Temperature scaling maps for each

corresponding flanking sequence size, illustrating how raw predicted probabilities for acceptor (x-axis) and donor (y-axis) sites are transformed after calibration. Arrows indicate the shift from pre- to post-calibration states in two-dimensional probability space.

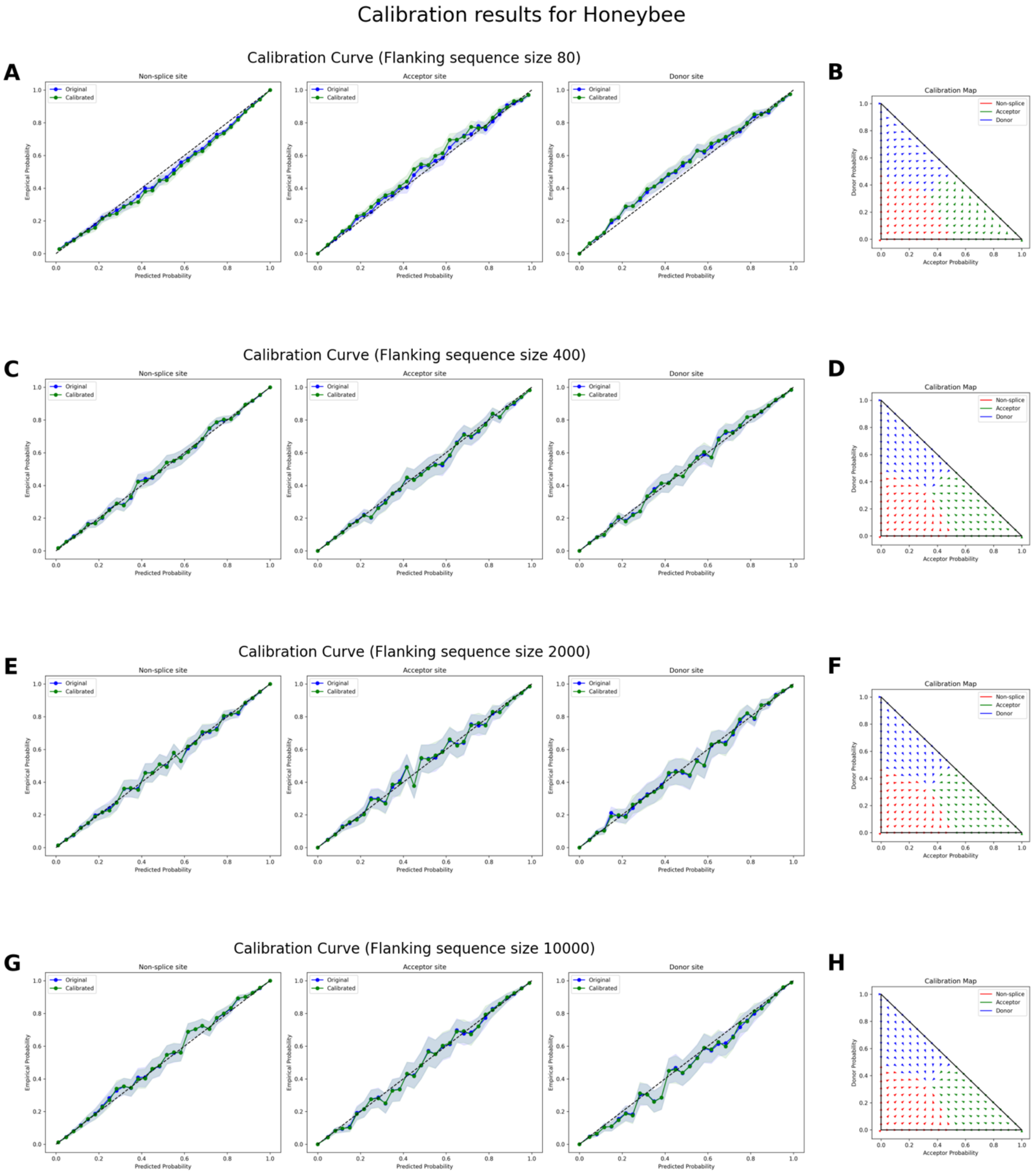


**Figure S21.**. Calibration results for honeybee (*Apis mellifera*) splice site classification at four flanking sequence sizes. **(A, C, E, G)** Reliability (calibration) curves for flanking sequence sizes of 80, 400, 2000, and 10,000 nucleotides, respectively. Each plot compares predicted probabilities (x-axis) to empirical probabilities (y-axis) for non-splice sites (left), acceptor sites (middle), and donor sites (right). The blue curves depict the reliability of the original $\text{OSAI}_{\text{Honeybee}}$ models, while the green curves show reliability after calibration. Shaded regions represent confidence intervals. The diagonal black line indicates perfect calibration, where predicted probabilities match observed frequencies exactly. **(B, D, F, H)** Temperature scaling maps for each corresponding flanking sequence size, illustrating how raw predicted probabilities for acceptor (x-axis) and donor (y-axis) sites are transformed after calibration. Arrows indicate the shift from pre- to post-calibration states in two-dimensional probability space.

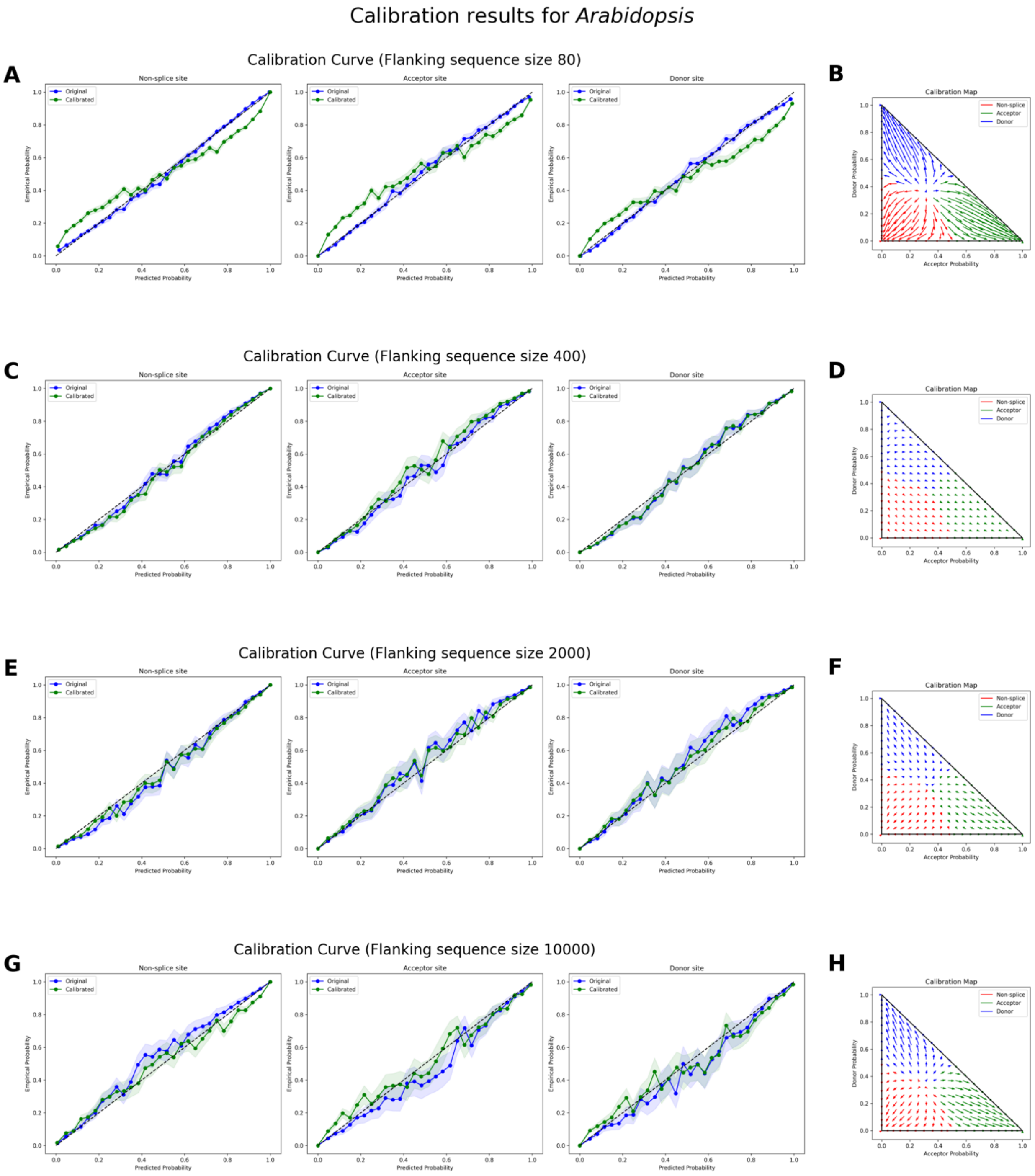


**Figure S22.**. Calibration results for *Arabidopsis thaliana* splice site classification at four flanking sequence sizes. **(A, C, E, G)** Reliability (calibration) curves for flanking sequence sizes of 80, 400, 2000, and 10,000 nucleotides, respectively. Each plot compares predicted probabilities (x-axis) to empirical probabilities (y-axis) for non-splice sites (left), acceptor sites (middle), and donor sites (right). The blue curves depict the reliability of the original $OSAI_{Arabidopsis}$ models, while the green curves show reliability after calibration. Shaded regions represent confidence intervals. The diagonal black line indicates perfect calibration, where predicted probabilities match observed frequencies exactly. **(B, D, F, H)** Temperature scaling maps for each

corresponding flanking sequence size, illustrating how raw predicted probabilities for acceptor (x-axis) and donor (y-axis) sites are transformed after calibration. Arrows indicate the shift from pre- to post-calibration states in two-dimensional probability space.

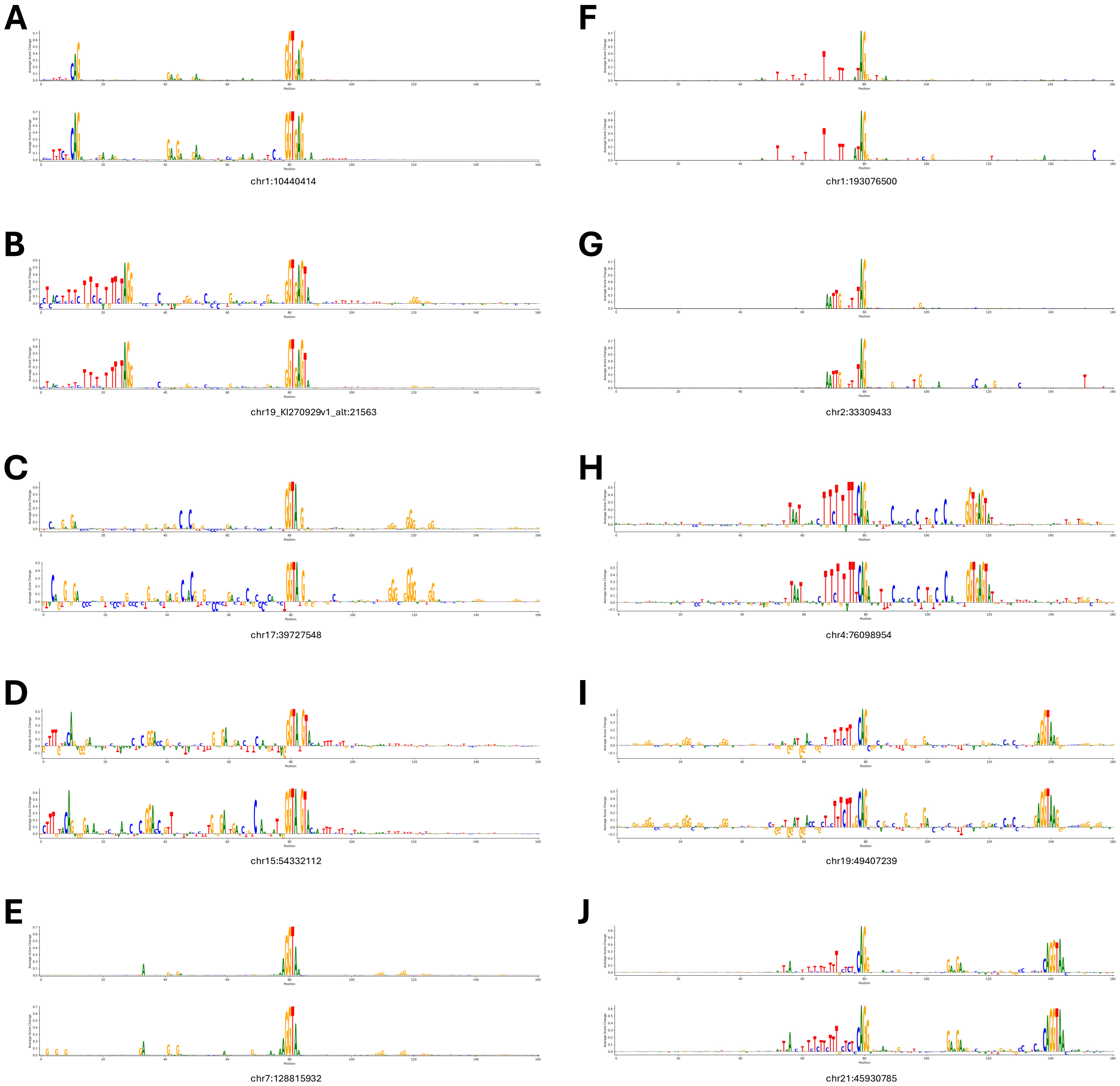


**Figure S23.** Zoomed-in 160 bp DNA sequence logos derived from ISM importance-score profiles for representative donor **(A–E)** and acceptor **(F–J)** splice sites. Logos were generated by mapping the ISM importance score at each position centered on the splice site to letter heights, with SpliceAI scores shown in the upper logo and $\text{OSAI}_{\text{MANE}}$ scores shown in the lower logo of each panel.

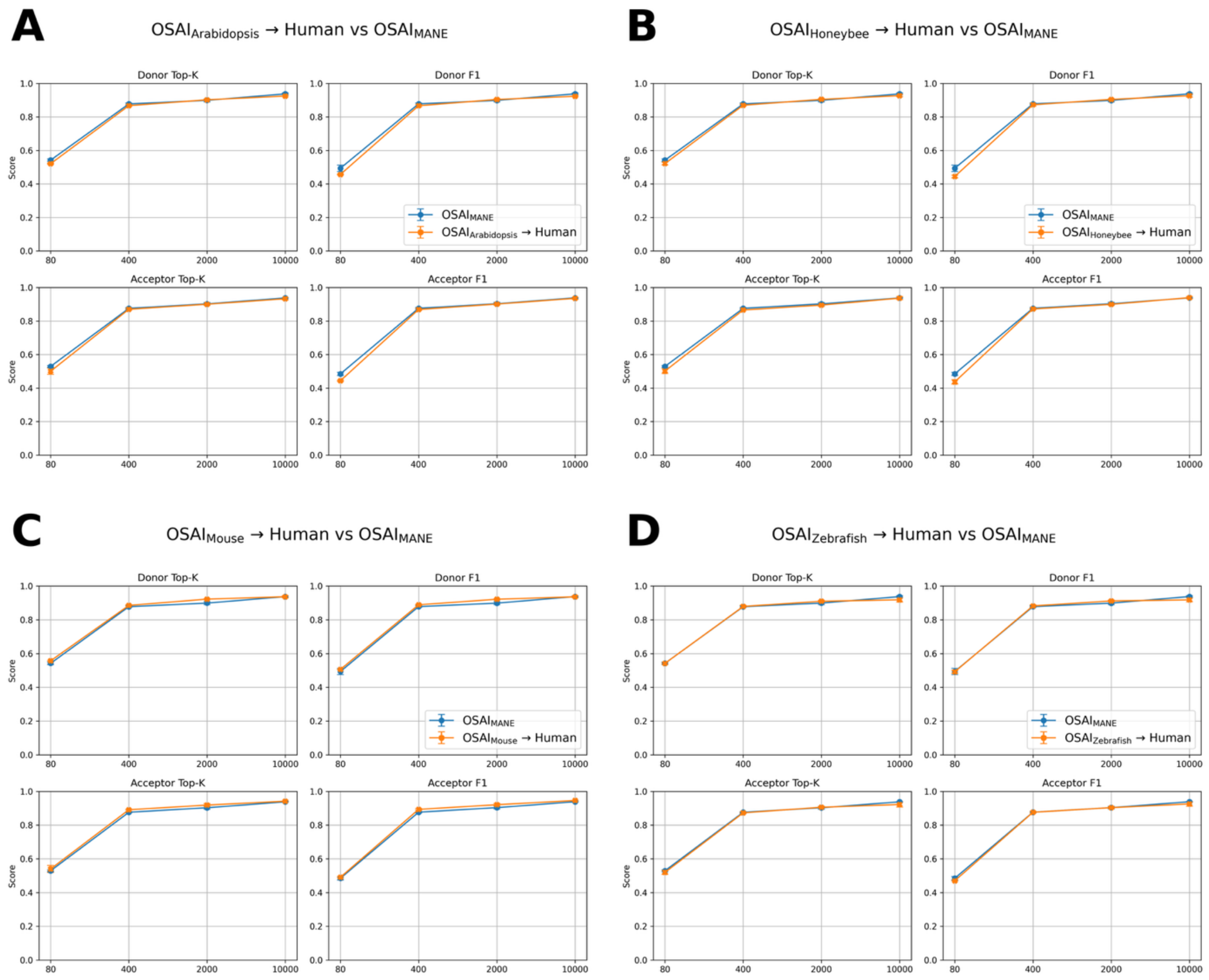


**Figure S24.** Cross-species transfer learning performance on human splice-site prediction. Models pretrained on **(A)** *Arabidopsis thaliana* ($\text{OSAI}_{\text{Arabidopsis}}$), **(B)** *Apis mellifera* ($\text{OSAI}_{\text{Honeybee}}$), **(C)** *Mus musculus* ($\text{OSAI}_{\text{Mouse}}$), and **(D)** *Danio rerio* ($\text{OSAI}_{\text{Zebrafish}}$) were fine-tuned on the human MANE dataset (orange) and compared to a model trained from scratch on MANE ($\text{OSAI}_{\text{MANE}}$, blue). Within each panel, the top row shows donor Top-K (left) and donor F1 (right) scores, and the bottom row shows acceptor Top-K (left) and acceptor F1 (right) scores. The x-axis indicates models trained with flanking sequence sizes of 80, 400, 2,000, and 10,000 bp.

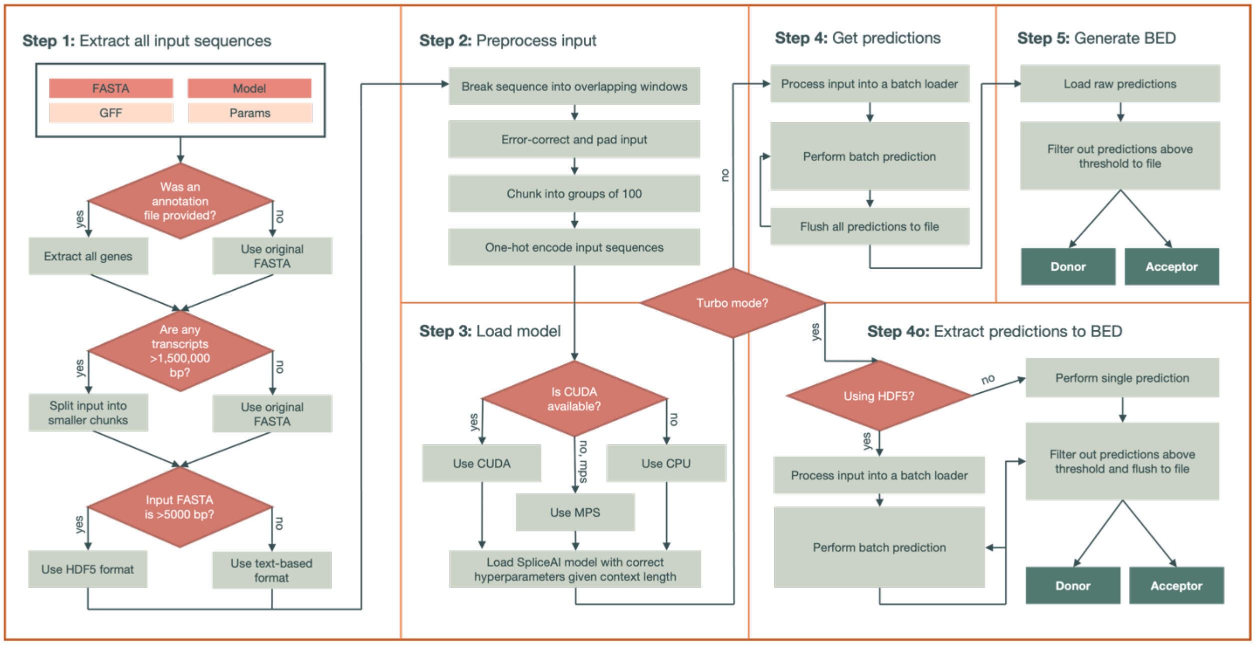


**Figure S25.** Decision-making and workflow of the predict subcommand. Required inputs include the FASTA file and PyTorch model, while optional inputs include a GFF annotation file and custom parameters. The outputs are two BED files corresponding to predicted donor and acceptor splice sites. Several intermediate files may be generated and useful to the user, including the HDF5-compressed datafile (raw sequences) and dataset (encoded inputs) for training.

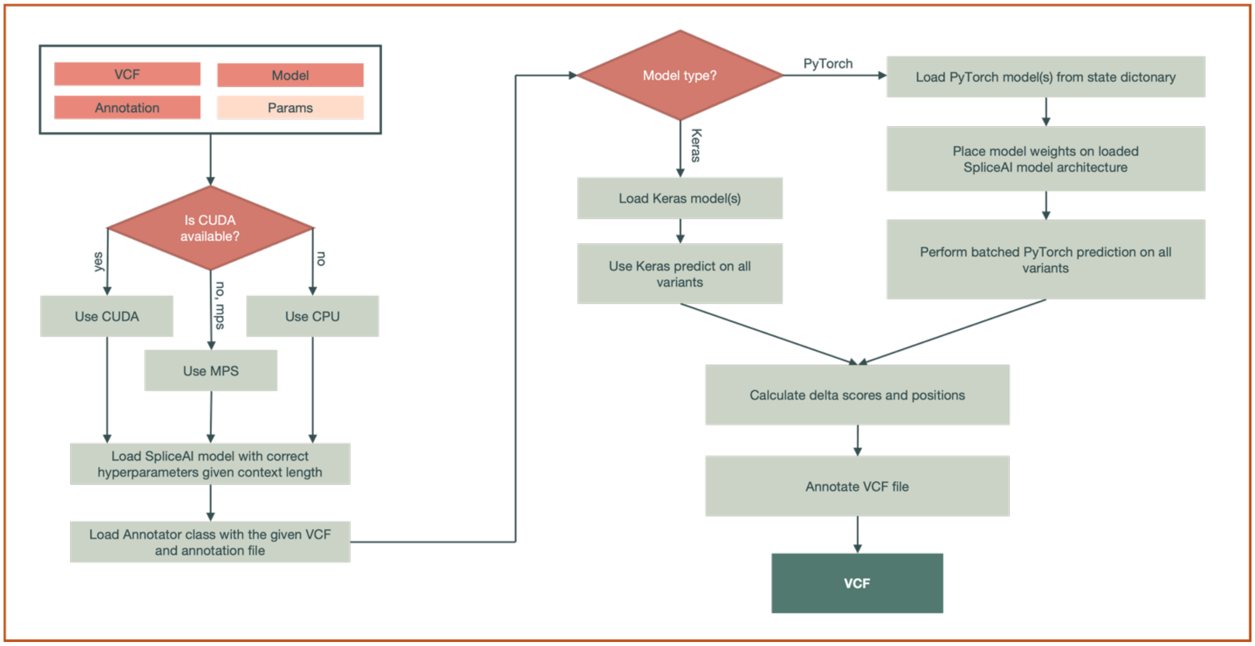


**Figure S26.** Decision-making and workflow of the variant subcommand. Required inputs include the VCF file, FASTA file, and annotation file, while optional inputs include the output path, splicing model, and distance, precision, and masking parameters. The output is a VCF file with the OpenSpliceAI delta scores of the variants. Note that both Keras and PyTorch models are supported in this tool, but PyTorch is recommended.

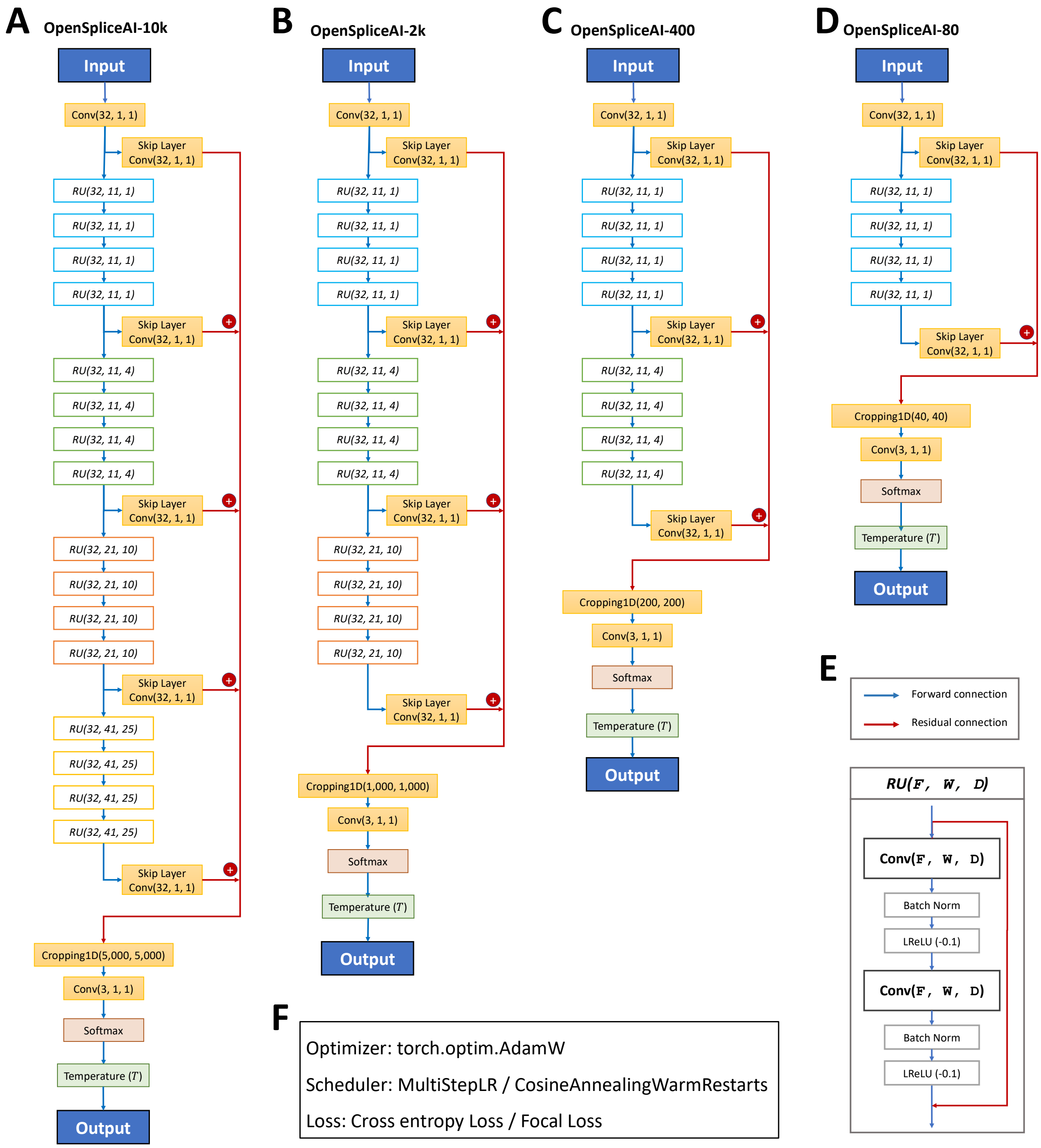


**Figure S27.** Overview of the OpenSpliceAI architectures trained with different flanking sequence lengths. **(A)** OpenSpliceAI-10k: Schematic of the model configured for 10 kb flanking regions. The input sequence is passed through an initial convolution layer (Conv1D), followed by 16 residual units, each incorporating skip connections. The final output is fed into a softmax function for splice site classification. **(B)** OpenSpliceAI-2k: Model variant with 2 kb flanking sequences, using a similar structure with 12 repeated residual units. **(C)** OpenSpliceAI-400: Model variant with 400 bp flanking sequences, using a similar structure with 8 repeated residual units. **(D)** OpenSpliceAI-80: The smallest variant, trained on 80 bp flanking sequences, using a similar structure with 4 repeated residual units. **(E)** Detailed view of the residual unit (RU) structure, highlighting the convolution, batch normalization, and skip connections. **(F)** Training setup: All models were trained using the AdamW optimizer, either a MultiStepLR or CosineAnnealingWarmRestarts learning rate scheduler, and cross-entropy or focal loss functions.
