## Supplemental Notes for "OpenSpliceAI: An efficient, modular implementation of SpliceAI enabling easy retraining on non-human species"

[†]These authors contributed equally

**Note S1.** Commands to train $\text{OSAI}_{\text{MANE}}$

- create-data

```
opensplicea create-data \
    --remove-paralogs \
    --min-identity 0.8 \
    --min-coverage 0.8 \
    --genome-fasta GCF_000001405.40_GRCh38.p14_genomic.fna \
    --annotation-gff MANE.GRCh38.v1.3.refseq_genomic.gff \
    --output-dir train_test_dataset_MANE/ \
    --parse-type canonical \
    --write-fasta \
    --split-method human --canonical-only >
train_test_dataset_MANE/output.log \
    2> train_test_dataset_MANE/error.log
```

- train

```
openspliceai train --flanking-size 10000 \
  --train-dataset train_test_dataset_MANE/dataset_train.h5 \
  --test-dataset train_test_dataset_MANE/dataset_test.h5 \
  --output-dir train_outdir/MANE/flanking_10000 \
  --project-name MANE_train \
  --exp-num 1 \
  --random-seed 10 \
  --epochs 10 \
  --scheduler CosineAnnealingWarmRestarts \
  --loss cross_entropy_loss >
train_outdir/MANE/flanking_10000/SpliceAI_MANE_train_10000_1_rs10/output.lo
g \
  2>
train_outdir/MANE/flanking_10000/SpliceAI_MANE_train_10000_1_rs10/error.log
```

- calibrate

```
openspliceai calibrate --flanking-size 10000 \
  --train-dataset train_test_dataset_MANE/dataset_train.h5 \
  --test-dataset train_test_dataset_MANE/dataset_test.h5 \
  --output-dir calibrate_outdir/MANE/flanking_10000 \
  --project-name human_MANE_calibrate \
  --random-seed 10 \
  --pretrained-model OSAI_MANE_model_10000nt_rs10.pt \
  --loss cross_entropy_loss >
calibrate_outdir/MANE/flanking_10000/output.log \
  2> calibrate_outdir/MANE/flanking_10000/error.log
```

**Note S2.** Commands to train OSAI$_{Mouse}$,

- create-data

```
openspliceai create-data \
    --remove-paralogs \
    --min-identity 0.8 \
    --min-coverage 0.8 \
    --genome-fasta GCF_000001635.27_GRCm39_genomic.fna \
    --annotation-gff GCF_000001635.27_GRCm39_genomic.gff \
    --output-dir train_test_dataset_mouse/ \
    --parse-type canonical \
    --write-fasta > train_test_dataset_mouse/output.log \
    2> train_test_dataset_mouse/error.log
```

- train

```
openspliceai train --flanking-size 10000 \
  --train-dataset train_test_dataset_mouse/dataset_train.h5 \
  --test-dataset train_test_dataset_mouse/dataset_test.h5 \
  --output-dir train_outdir/mouse/flanking_10000 \
  --project-name mouse_train \
  --exp-num 1 \
  --random-seed 10 \
  --epochs 10 \
  --scheduler CosineAnnealingWarmRestarts \
  --loss cross_entropy_loss >
train_outdir/mouse/flanking_10000/SpliceAI_mouse_train_10000_1_rs10/output.
log \
  2>
train_outdir/mouse/flanking_10000/SpliceAI_mouse_train_10000_1_rs10/error.l
og
```

- calibrate

```
openspliceai calibrate --flanking-size 10000 \

  --train-dataset train_test_dataset_mouse/dataset_train.h5 \
  --test-dataset train_test_dataset_mouse/dataset_test.h5 \
  --output-dir calibrate_outdir/mouse/flanking_10000 \
  --project-name human_mouse_calibrate \
  --random-seed 10 \
  --pretrained-model OSAI_mouse_model_10000nt_rs10.pt \
  --loss cross_entropy_loss >
calibrate_outdir/mouse/flanking_10000/output.log \
  2> calibrate_outdir/mouse/flanking_10000/error.log
```

- transfer

```
openspliceai fine-tune --flanking-size 10000 \
  --train-dataset train_test_dataset_mouse/dataset_train.h5 \
  --test-dataset train_test_dataset_mouse/dataset_test.h5 \
  --output-dir transfer_outdir/mouse/flanking_10000 \
  --project-name human_mouse_fine-tune \
  --exp-num 0 \
  --random-seed 10 \
```

```
  --pretrained-model OSAI_MANE_model_10000nt_rs10.pth \
  --epochs 10 \
  --scheduler CosineAnnealingWarmRestarts \
  --loss cross_entropy_loss >
transfer_outdir/mouse/flanking_10000/SpliceAI_human_mouse_transfer_10000_0_
rs10/output.log \
  2>
transfer_outdir/mouse/flanking_10000/SpliceAI_human_mouse_transfer_10000_0_
rs10/error.log
```

**Note S3.** Commands to train $OSAI_{Zebrafish}$

- create-data

```
openspliceai create-data \
    --remove-paralogs \
    --min-identity 0.8 \
    --min-coverage 0.8 \
    --genome-fasta GCF_000002035.6_GRCz11_genomic.fna \
    --annotation-gff  GCF_000002035.6_GRCz11_genomic.gff \
    --output-dir train_test_dataset_zebrafish/ \
    --parse-type canonical \
    --write-fasta > train_test_dataset_zebrafish/output.log \
    2> train_test_dataset_zebrafish/error.log
```

- train

```
openspliceai train --flanking-size 10000 \
  --train-dataset train_test_dataset_zebrafish/dataset_train.h5 \
  --test-dataset train_test_dataset_zebrafish/dataset_test.h5 \
  --output-dir train_outdir/zebrafish/flanking_10000 \
  --project-name zebrafish_train \
  --exp-num 1 \
  --random-seed 10 \
  --epochs 10 \
  --scheduler CosineAnnealingWarmRestarts \
  --loss cross_entropy_loss >
train_outdir/zebrafish/flanking_10000/SpliceAI_zebrafish_train_10000_1_rs10
/output.log \
  2>
train_outdir/zebrafish/flanking_10000/SpliceAI_zebrafish_train_10000_1_rs10
/error.log
```

- calibrate

```
openspliceai calibrate --flanking-size 10000 \
  --train-dataset train_test_dataset_zebrafish/dataset_train.h5 \
  --test-dataset train_test_dataset_zebrafish/dataset_test.h5 \
  --output-dir calibrate_outdir/zebrafish/flanking_10000 \
  --project-name human_zebrafish_calibrate \
  --random-seed 10 \
  --pretrained-model OSAI_zebrafish_model_10000nt_rs10.pt \
  --loss cross_entropy_loss >
calibrate_outdir/zebrafish/flanking_10000/output.log \
  2> calibrate_outdir/zebrafish/flanking_10000/error.log
```

- transfer

```
openspliceai fine-tune --flanking-size 10000 \
  --train-dataset train_test_dataset_zebrafish/dataset_train.h5 \
  --test-dataset train_test_dataset_zebrafish/dataset_test.h5 \
  --output-dir transfer_outdir/zebrafish/flanking_10000 \
  --project-name human_zebrafish_fine-tune \
  --exp-num 0 \
```

```
  --random-seed 10 \
  --pretrained-model OSAI_MANE_model_10000nt_rs10.pth \
  --epochs 10 \
  --scheduler CosineAnnealingWarmRestarts \
  --loss cross_entropy_loss >
transfer_outdir/zebrafish/flanking_10000/SpliceAI_human_zebrafish_transfer_
10000_0_rs10/output.log \
  2>
transfer_outdir/zebrafish/flanking_10000/SpliceAI_human_zebrafish_transfer_
10000_0_rs10/error.log
```

**Note S4.** $\text{OSAI}_{\text{Honeybee}}$

- create-data

```
openspliceai create-data \
    --remove-paralogs \
    --min-identity 0.8 \
    --min-coverage 0.8 \
    --genome-fasta HAv3.1_genomic.fna \
    --annotation-gff HAv3.1_genomic.gff \
    --output-dir train_test_dataset_honeybee/ \
    --parse-type canonical \
    --write-fasta > train_test_dataset_honeybee/output.log \
    2> train_test_dataset_honeybee/error.log
```

- train

```
openspliceai train --flanking-size 10000 \
  --train-dataset train_test_dataset_honeybee/dataset_train.h5 \
  --test-dataset train_test_dataset_honeybee/dataset_test.h5 \
  --output-dir train_outdir/honeybee/flanking_10000 \
  --project-name honeybee_train \
  --exp-num 1 \
  --random-seed 10 \
  --epochs 10 \
  --scheduler CosineAnnealingWarmRestarts \
  --loss cross_entropy_loss >
train_outdir/honeybee/flanking_10000/SpliceAI_honeybee_train_10000_1_rs10/o
utput.log \
  2>
train_outdir/honeybee/flanking_10000/SpliceAI_honeybee_train_10000_1_rs10/e
rror.log
```

- calibrate

```
openspliceai calibrate --flanking-size 10000 \
  --train-dataset train_test_dataset_honeybee/dataset_train.h5 \
  --test-dataset train_test_dataset_honeybee/dataset_test.h5 \
  --output-dir calibrate_outdir/honeybee/flanking_10000 \
  --project-name human_honeybee_calibrate \
  --random-seed 10 \
  --pretrained-model OSAI_honeybee_model_10000nt_rs10.pt \
  --loss cross_entropy_loss >
calibrate_outdir/honeybee/flanking_10000/output.log \
  2> calibrate_outdir/honeybee/flanking_10000/error.log
```

- transfer

```
openspliceai fine-tune --flanking-size 10000 \
  --train-dataset train_test_dataset_honeybee/dataset_train.h5 \
  --test-dataset train_test_dataset_honeybee/dataset_test.h5 \
  --output-dir transfer_outdir/honeybee/flanking_10000 \
  --project-name human_honeybee_fine-tune \
  --exp-num 0 \
  --random-seed 10 \
```

```
  --pretrained-model OSAI_MANE_model_10000nt_rs10.pth \
  --epochs 10 \
  --scheduler CosineAnnealingWarmRestarts \
  --loss cross_entropy_loss >
transfer_outdir/honeybee/flanking_10000/SpliceAI_human_honeybee_transfer_10
000_0_rs10/output.log \
  2>
transfer_outdir/honeybee/flanking_10000/SpliceAI_human_honeybee_transfer_10
000_0_rs10/error.log
```

**Note S5.** $OSAI_{Arabidopsis}$

- create-data

```
openspliceai create-data \
    --remove-paralogs \
    --min-identity 0.8 \
    --min-coverage 0.8 \
    --genome-fasta TAIR10.fna \
    --annotation-gff TAIR10.gff \
    --output-dir train_test_dataset_arabidopsis/ \
    --parse-type canonical \
    --write-fasta > train_test_dataset_arabidopsis/output.log \
    2> train_test_dataset_arabidopsis/error.log
```

- train

```
openspliceai train --flanking-size 10000 \
    --train-dataset train_test_dataset_arabidopsis/dataset_train.h5 \
    --test-dataset train_test_dataset_arabidopsis/dataset_test.h5 \
    --output-dir train_outdir/arabidopsis/flanking_10000 \
    --project-name arabidopsis_train \
    --exp-num 20 \
    --random-seed 10 \
    --epochs 10 \
    --scheduler CosineAnnealingWarmRestarts \
    --loss cross_entropy_loss >
train_outdir/arabidopsis/flanking_10000/SpliceAI_arabidopsis_train_10000_20
_rs10/output.log 2>
train_outdir/arabidopsis/flanking_10000/SpliceAI_arabidopsis_train_10000_20
_rs10/error.log
```

- calibrate

```
openspliceai calibrate --flanking-size 10000 \
  --train-dataset train_test_dataset_arabidopsis/dataset_train.h5 \
  --test-dataset train_test_dataset_arabidopsis/dataset_test.h5 \
  --output-dir calibrate_outdir/arabidopsis/flanking_10000 \
  --project-name human_arabidopsis_calibrate \
  --random-seed 10 \
  --pretrained-model OSAI_arabidopsis_model_10000nt_rs10.pt \
  --loss cross_entropy_loss >
calibrate_outdir/arabidopsis/flanking_10000/output.log \
  2> calibrate_outdir/arabidopsis/flanking_10000/error.log
```

- transfer

```
openspliceai fine-tune --flanking-size 10000 \
  --train-dataset train_test_dataset_arabidopsis/dataset_train.h5 \
  --test-dataset train_test_dataset_arabidopsis/dataset_test.h5 \
  --output-dir transfer_outdir/arabidopsis/flanking_10000 \
  --project-name human_arabidopsis_fine-tune \
  --exp-num 0 \
  --random-seed 10 \
  --pretrained-model OSAI_MANE_model_10000nt_rs10.pth \
```

```
  --epochs 10 \
  --scheduler CosineAnnealingWarmRestarts \
  --loss cross_entropy_loss >
transfer_outdir/arabidopsis/flanking_10000/SpliceAI_human_arabidopsis_trans
fer_10000_0_rs10/output.log \
  2>
transfer_outdir/arabidopsis/flanking_10000/SpliceAI_human_arabidopsis_trans
fer_10000_0_rs10/error.log
```

### Note S6. Discussion of the enhancements in GPU memory management and efficiency in OpenSpliceAI

A key improvement of OpenSpliceAI over the original SpliceAI is its optimized approach to GPU memory management – a critical factor for handling large-scale genomic datasets. The original SpliceAI, built on Keras with TensorFlow, relies on a static computation graph that requires pre-allocation of memory based on the worst-case input size. This design forces the model to reserve large memory buffers, leading to inefficient memory utilization and an increased risk of out-of-memory (OOM) errors. Furthermore, TensorFlow's default behavior aggressively pre-allocates GPU memory – even when it is not fully utilized – which limits resource availability for concurrent tasks or for deploying multiple models (for example, five models are used per flanking sequence). In contrast, OpenSpliceAI leverages PyTorch's dynamic computation graph and memory allocator, which allocate GPU memory on demand and efficiently cache, release, and reuse memory. This integrated approach not only reduces the overall memory footprint but also enhances flexibility and robustness for both DNA sequence predictions and genetic variant splice site loss/gain predictions.

Batch processing is also significantly enhanced in OpenSpliceAI. The original SpliceAI does not implement batch prediction for its trained models, requiring users to process each input sequence sequentially. When users iterate over sequences of varying lengths in a large loop, it results in GPU context-switching overhead, and the pre-allocated memory is not properly released, leading to out-of-memory (OOM) errors. In contrast, OpenSpliceAI implements batch prediction. For example, the OpenSpliceAI-10k model segments the input sequence into chunks of 5,000 base pairs, with an additional 5,000 base pairs of flanking sequence on each side, using a stride of 5,000. Although gene sequences are segmented for tensor conversion during batch prediction, this approach ensures every output position fully leverages the model's receptive field, thereby optimizing performance and memory usage.

In summary, the transition from a Keras/TensorFlow-based implementation to one built on PyTorch has enabled OpenSpliceAI to overcome the inherent limitations of static graph-based GPU memory management. By adopting dynamic memory allocation, and implementing batch prediction and parallelization, OpenSpliceAI achieves lower memory usage, reduces the frequency of OOM errors, and improves overall processing efficiency. These improvements make OpenSpliceAI a more robust and scalable tool and enable more efficient use of GPU resources not only in the gene-level, but also genome-level prediction.
